# Balance between inhibitory cell types is necessary for flexible frequency switching in adult mouse visual cortex

**DOI:** 10.1101/2020.01.18.911271

**Authors:** Justin W. M. Domhof, Paul H. E. Tiesinga

**Affiliations:** Donders Centre for Neuroscience, Radboud University, Nijmegen, The Netherlands

## Abstract

Neuronal networks in rodent primary visual cortex (V1) can generate oscillations in different frequency bands depending on the network state and the level of visual stimulation. High-frequency gamma rhythms, for example, dominate the network’s spontaneous activity in adult mice but are attenuated upon visual stimulation, during which the network switches to the beta band instead. The spontaneous LFP of juvenile mouse V1, however, mainly contains beta oscillations and presenting a stimulus does not elicit drastic changes in collective network oscillations. We study, in a spiking neuron network model, the mechanism in adult mice that allows for flexible switches between multiple frequency bands and contrast this to the network structure in juvenile mice that do not posses this flexibility. The model is comprised of excitatory pyramidal cells (PCs) and two types of inhibitory interneurons: the parvalbumin expressing (PV) interneuron, which produces gamma oscillations, and the somatostatin expressing (SOM) cell, which generates beta rhythms. Our model simulations suggest that both of these oscillations are generated by a pyramidal-interneuron gamma (PING) mechanism. Furthermore, prominent gamma and beta oscillations in, respectively, the spontaneous and visually evoked activity of the simulated network only occurred within the same network configuration when there was a balance between both types of interneurons so that SOM neurons are able to shape the dynamics of the pyramidal-PV cell subnetwork without dominating dynamics. Taken together, our results demonstrate that the effective strengths of PV and SOM cells must be balanced for experimentally observed V1 dynamics in adult mice. Moreover, since spontaneous gamma rhythms emerge during the well-known critical period, our findings support the notion that PV cells become integrated in the circuit of this cortical area during this time window and additionally indicate that this integration comprises an overall increase in their synaptic strength.

## Introduction

Over the past decade, the primary visual cortex (V1) of mice has been intensively studied using transgenic techniques. Distinct subtypes of inhibitory interneurons that use gamma-aminobutyric acid (GABA) as neuro-transmitter were, for example, identified and linked to various spiking behaviors [1, 2]. These different subpopulations also have specific connectivity patterns with respect to one another; parvalbumin-expressing (PV) interneurons are mutually inhibited and receive inhibition from somatostatin-expressing (SOM) cells, while the latter subtype primarily receives inhibition from vasoactive intestinal peptide expressing (VIP) interneurons [3]. These subtype-specific connection motifs also have an additional geometric component: PV interneurons in cortical layer 2 and 3 (L2/3) predominantly receive vertical inputs originating from cortical layer 4 (L4) compared to horizontal ones from other L2/3 cells, whereas the opposite is true for SOM cells as these receive more horizontally than vertically aligned inputs [4].

This particular subtype-specific arrangement of the a- and efferents within the network probably underlies the different synchronization regimes that have been observed in mouse V1 [5, 6, 7]. One of these changes is the attenuation of gamma oscillations when visually stimulating the adult mouse; synchronization is instead achieved in the beta frequency band (Figure 1C–D) [5, 6]. The extent to which gamma and beta rhythms are decreased and enhanced, respectively, depends on the size of the stimulus, which led to the hypothesis that an inhibitory surround effect underlies this switch in synchrony [4, 6]. These gamma oscillations are not present in younger mice as they develop during the critical period (CP) (Figure 1A), a period well-known for its increased levels of ocular dominance plasticity [5, 8]. Furthermore, before the CP, visual stimulation does not alter the network’s synchronization frequency, even though beta power is still slightly enhanced (Figure 1B) [5]. What exactly initiates the development of gamma rhythms during this time window is unclear, but PV cells are likely involved: studies using optogenetic perturbations indicate that PV and SOM cells can be associated with the gamma and beta oscillations, respectively [6, 9, 10]. PV cell progenitors transplanted into the V1 of adult mice differentiate into the PV (hence not the SOM) subtype, integrate successfully in the network and initiate a period with enhanced plasticity that resembles the CP [11, 12]. These findings thus indicate that PV cells are involved in gamma oscillation generation and in CP associated plasticity, but even more evidence can be found in the experimental literature that strengths these associations. One other study, for instance, shows that gamma rhythms in V1 are transiently amplified right after the start of monocular deprivation (MD) in juvenile mice and in adult ones that have been depleted of the perineuronal nets (PNNs), which pre-dominantly ensheath PV cells, in that area [7, 13]. Although the exact cause for this rise in gamma power is unknown, it has been shown that, in the same time interval after the start of MD during the CP, the spike rates of pyramidal cells (PCs) were at first reduced but gradually rose to their original values through a decrease of the PV cell activity in the network [14]. Taken together, these two findings also support the notion that PV cells generate gamma rhythms and are involved in the plasticity mechanisms that are active during the CP. With respect to the former of these two studies, however, it must be mentioned that there is a controversy as to whether the stronger gamma oscillations following PNN removal critically depend on MD. A recent study, namely, has demonstrated that merely removing the PNNs results in enhanced gamma power as well and that stronger thalamic inputs to the PV cells in L4 could account for this; however, this rise in gamma power is attenuated by depleting the PNNs and starting MD immediately thereafter [15].

**Figure 1:**
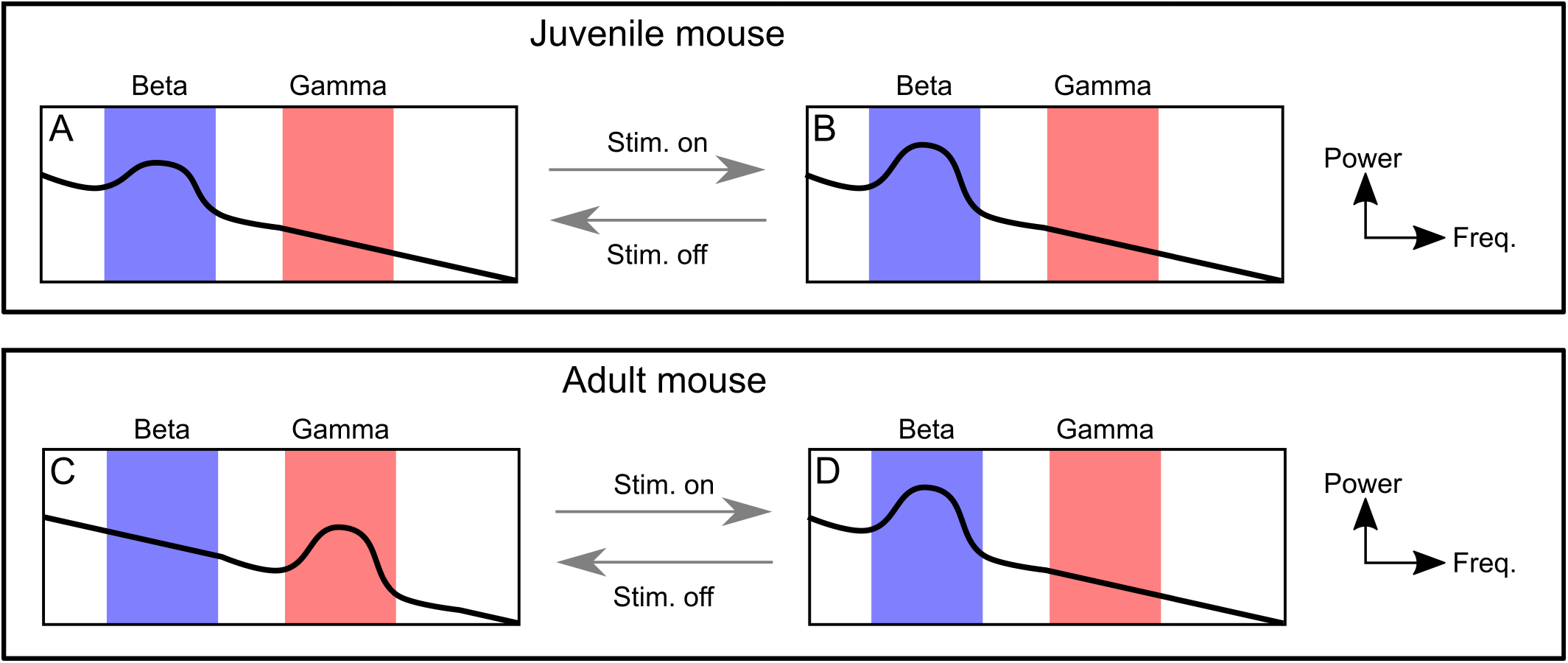
Visual stimulation elicits a change in the synchronization of the neural ensemble in the adult, but not juvenile, mouse primary visual cortex (V1) (A–B) Spectral density characterization of the spontaneous (A) and visually evoked (B) local field potential (LFP) in V1 of juvenile mice (< *P*20). (C–D) Spectral density characterization of the spontaneous (C) and visually evoked (D) LFP in adult mouse V1 (> *P*60). Abbreviations: stim. = stimulus.

So far, the phenomenon of surround inhibition and the shift from gamma to beta oscillations via visual stimulation has only been explained at the level of neural mass models [6, 16]. These models, however, do not reveal the spiking-dependent network mechanisms and, furthermore, do not explain the emergence of gamma oscillations during the CP. Therefore, we, inspired by the phenomena mentioned above, developed a spiking neuron network model comprising pyramidal, PV and SOM cells to determine the network configurations that enable the switch in synchronization from the gamma to the beta frequency band upon visual stimulation.

Separate considerations of the pyramidal–PV and pyramidal–SOM cell subnetworks in our model not only confirm the hypotheses that PV and SOM cells are involved in the generation of gamma and beta oscillations respectively, but additionally suggest that these rhythms are both realized via pyramidal-interneuron gamma (PING) mechanisms. Analogously, the results acquired through the variation of the size of the stimulated area agree with previous models that the respective enhancement and attenuation of beta and gamma rhythms through visual stimulation are a consequence of surround inhibition and, in addition, show that this switch is realized by SOM cells outcompeting the PV cells generating the gamma oscillation so that the network produces beta rhythms instead. Finally, by sampling network activity for various settings of the PV and SOM cell projection strength, we demonstrate that only a restricted range of values for these parameters gives rise to the attenuation of gamma and amplification of beta oscillations following visual stimulation. This indicates that these two subtypes must exert an approximately equal influence on the PCs in adult mice; if this balance is not present, the characteristic switch in the network synchronization frequency cannot be induced in one and the same network realization. Moreover, given that the spontaneous LFP of the pre-CP mouse V1 contains primarily beta oscillations and the post-CP one predominantly gamma, our result also suggest that PV cell projections are, on average, strengthened across the CP, which is consistent with the notion of PV cells being integrated in the local network of this cortical area during that time window.

## Method

The results presented in this article were obtained through means of a network model, which was comprised of 4500 neurons. These neurons were modeled using Izhikevich’s model [17] and received input from three sources: background activity, visually induced currents and recurrent inputs. With respect to the latter input source, the connectivity patterns depicted in Figure 2A were implemented. The model’s equations will be elaborated upon in the remainder of this section, followed by a brief description explaining how they were implemented and how the analysis was carried out.

**Figure 2:**
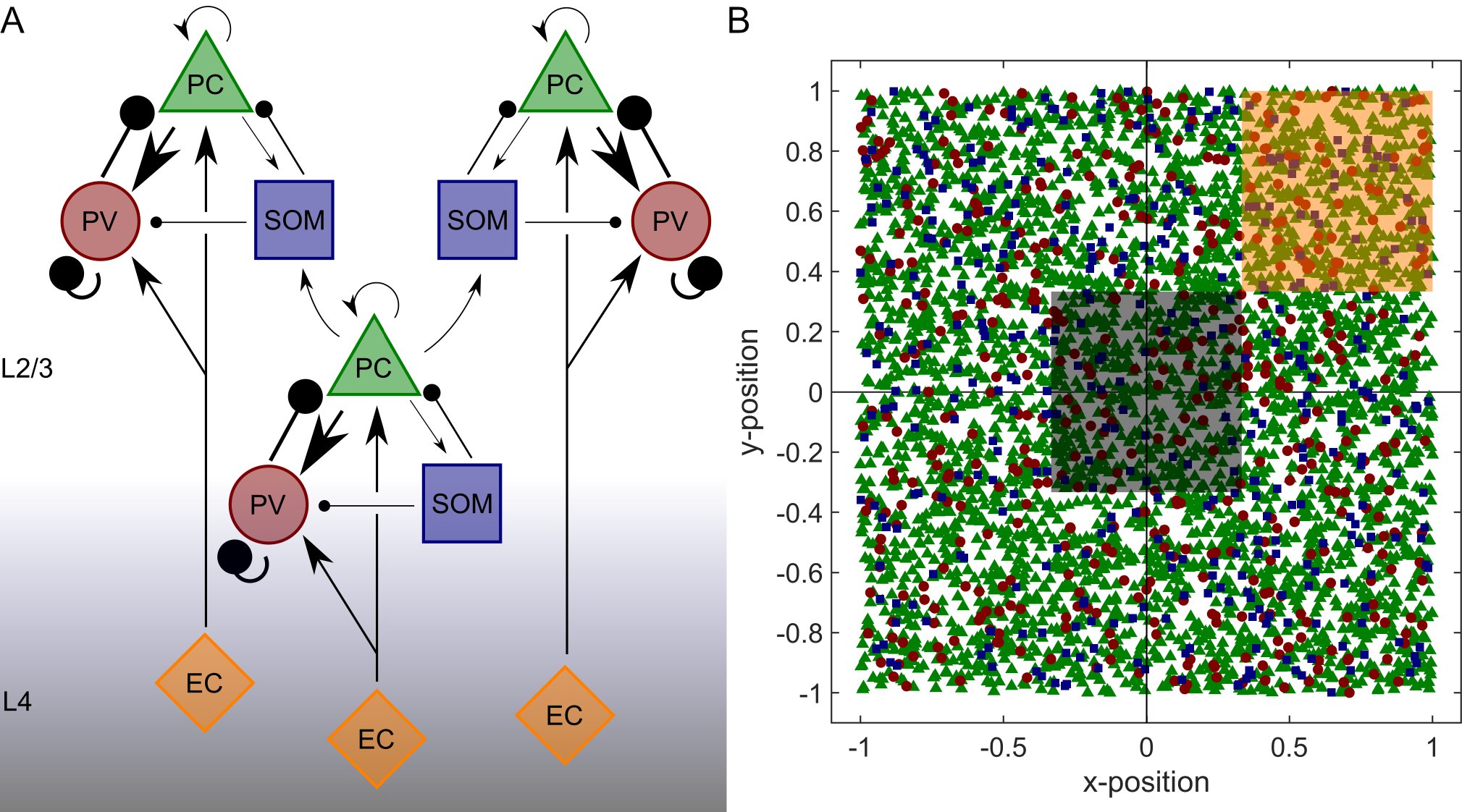
Schematic of the subtype-specific connectivity patterns in the network, cell positions and graphical depiction of areas used for the analysis. (A) Schematic that shows the connectivity patterns embedded in the network. Long-distance connections are only shown for the center column. (B) Example of cell positioning. Green triangles, red dots and blue squares represent pyramidal, parvalbumin expressing (PV) and somatostatin expressing (SOM) neurons, respectively. The black and orange shaded areas additionally depict the center and reference regions. Other abbreviations: EC = excitatory cell from cortical layer 4, L2/3 = cortical layer 2 and 3, L4 = cortical layer 4.

### Membrane potential dynamics

In our model, a total of 4500 neurons situated in L2/3 were considered: *N*^*PC*^ = 3600 pyramidal, *N*^*PV*^ = 495 PV and *N*^*SOM*^ = 405 SOM neurons, which roughly matched the reported relative abundances of these cell types in V1 [2, 18]. They were distributed across a 2 × 2 square patch by randomly assigning them positions in a Cartesian coordinate system 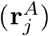 via

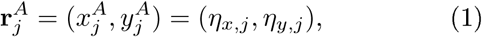

where the index *j* ∈ {1, 2, *…, N*^*A*^} denotes a cell within neuron population *A* ∈ {*P C, PV, SOM*} and *η*_*x,j*_ and *η*_*y,j*_ are independent random values drawn from a uniform distribution between −1 and 1. The square’s size was defined to be dimensionless as we wanted to investigate how the extent of the activated area relative to the entire patch of cortex influenced the oscillations generated by the model. An example of the resulting cell positions is shown in Figure 2B.

The dynamics of the membrane potentials were simulated using Izhikevich’s model, which comprises a set of two coupled differential equations [17]. The first of them specifies the membrane potential (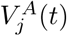, in Volts),

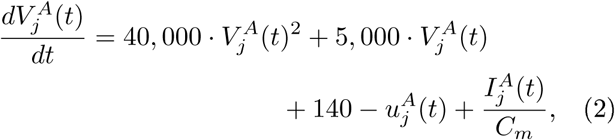

and the other describes a slower gating variable (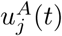, in Volts/second),

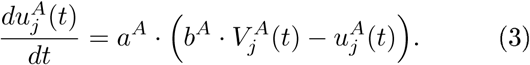

In these two equations, *C*_*m*_ and 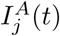 represent the membrane capacitance of and the total input current received by the neuron under consideration (see below), respectively. An action potential is detected when the membrane potential exceeds 30 mV; such a detection imposes the following reset condition on the membrane potential and the gating variable:

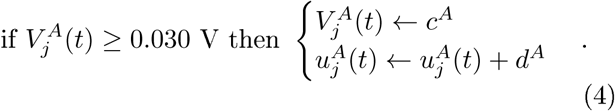

In the two preceding equations, *a*^*A*^, *b*^*A*^, *c*^*A*^ and *d*^*A*^ are parameters chosen such that the modeled spiking behavior resembles its experimentally observed counterpart. The parameter values are given in Table 1; these values were based on the values given in the literature for these cell types [1, 17], but have been adjusted in order to approximately match the oscillation frequencies emerging from the model to those observed in experimental studies [5, 6, 9]. Additionally, the cell capacitance was set to *C*_*m*_ = 100 pF for all subtypes.

**Table 1:** Values for the subtype-specific parameters in the model. A ± sign designates a Gaussian distributed random variable following the notation *mean* ± *standard deviation*. Abbreviations: PV = parvalbumin expressing, Pyr. = pyramidal, SOM = somatostatin expressing.

| Neuron type | a (s<sup>-1</sup>) | b (s<sup>-1</sup>) | c (mV) | d (V·s<sup>-1</sup>) | RK<sub>bg</sub><sup>A</sup> (Hz) | $\bar{g}_{bg}^A$ (pS·s) |
| --- | --- | --- | --- | --- | --- | --- |
| Pyr. | 40 ± 4 | 200 | -65.0 ± 6.5 | 8.0 ± 0.8 | 1,500 | 6 |
| PV | 100 ± 10 | 250 | -65.0 ± 6.5 | 2.0 ± 0.2 | 200 | 22 |
| SOM | 40 ± 4 | 250 | -65.0 ± 6.5 | 2.0 ± 0.2 | 1 | 6 |

We recognize that Eqs. 2–4 are in a somewhat different form than in [17]. In the Supplementary method it is explained how the forms shown here derive from our choice of units.

### Input currents

The total input current received by each cell consisted of three different components, which are the background currents 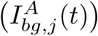, the visually induced input from L4 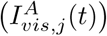 and the recurrent inputs 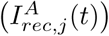:

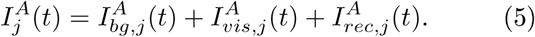

The background represents inputs originating from an ensemble of 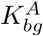 neurons that are not explicitly modeled but whose spike trains are characterized by Poisson processes with a constant firing rate of 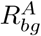. When the product of the latter two parameters is denoted by 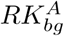, the resulting background current received by the neuron can be approximated via [19]

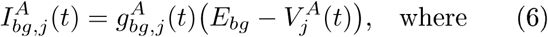

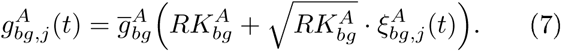

In these equations, 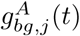 denotes the conductance induced in the cell and 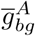 and *E*_*bg*_ are the overall scale of conductance induced by and the reversal potential of the background neuron ensemble, respectively. Finally, 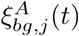 is a Gaussian distributed random variable with zero mean and a temporal correlation of 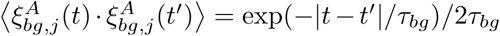. The values for the subtype-specific parameters are shown in Table 1. The remaining parameters, *E*_*bg*_ and *τ*_*bg*_, were respectively set to 0 mV and 2 ms. Finally, note that the value for 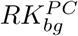 was changed in some simulations to assess whether a particular circuit produced rhythms via an interneuron gamma (ING) or a PING mechanism; if the excitability of the pyramidal cells is increased, an ING and PING mechanism give rise to similar and altered oscillation frequencies, respectively [20].

The dorsal part of the lateral geniculate nucleus, which receives its input from the retina, projects to L4 of V1 [21]. The signals are subsequently forwarded to L2/3 [22, 23]. Since the model represents L2/3 neurons, visually induced activity was therefore assumed to be provided by the afferents originating from the same horizontal location but in L4. Following a previous study [4], it was decided that only pyramidal and PV cells (*A* ∈ {*PC, PV*}) received visually induced currents that thus presumably originate from L4 (Figure 2A). Not all the cells within these subgroups received visual input: cells needed to be within a square field with length 2 · *D*_*vis*_ that was concentric with the square patch. Approximately half of the cells within this area were randomly selected to receive the visually induced currents. Taken together, the following condition thus had to be met by a pyramidal or PV cell to receive visually induced input:

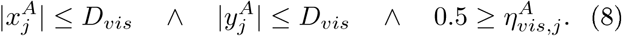

Here, ∧ is the logical AND operator and 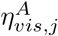 is a number randomly drawn from the standard uniform distribution (interval between 0 and 1). The visually induced current had the following form:

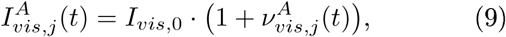

where *I*_*vis*,0_ represents the magnitude of the current and 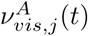 is Gaussian white noise with a mean of 0 and a standard deviation of 1. If not specified otherwise, *I*_*vis*,0_ had a value of 100 pA. The variable *D*_*vis*_ was varied to assess its effect on the network’s synchronization. It approximately reflected the effects of stimulus size, without specifically taking into account the retinotopic mapping as well as other feature maps that are present in the mouse visual cortex [24].

The recurrent inputs included the currents that the explicitly modeled neurons in the network received from one another. The probability of neuron *j* from subtype *A* to have neuron *k* from subtype *B* as an afferent was calculated via

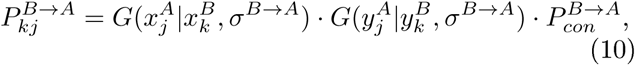

where the factor 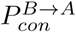 has been included so that the relative number of connections approximately matched the subtype-specific connection probabilities listed in the experimental literature [3, 25, 26]. In addition, the function *G*(*x*|*m, s*) represents the periodic Gaussian probability distribution function

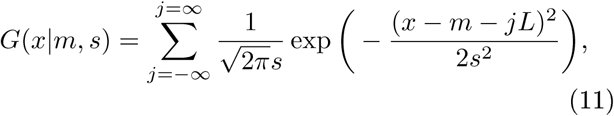

where *L* corresponds to the length of the square patch. In this context, one can interpret *σ*^*B*→*A*^ as the width of the area around a cell from subpopulation *B* wherein it is possible for that neuron to connect to a neuron from subtype *A*. Note also that Eq. 10 and 11 imply that the probability distribution is not strictly circularly symmetric. Instead, the periodic Gaussian increases the likelihood that a neuron close to one side of the square patch is connected to a cell that is near the opposite side so that the number of inputs to a neuron from a particular subtype is approximately equal throughout the square patch. The weights of the connections 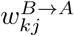 were initialized via

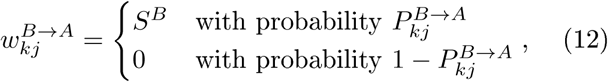

where *S*^*B*^ is the subtype-specific projection strength. Given these weights, the recurrent input received by neuron *j* from subpopulation *A* ∈{*PC, PV, SOM*} was determined via

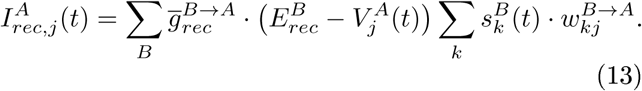

In this expression, 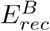 is the reversal potential of the synapse type associated with the cell type at the presynaptic side and 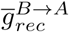 is the amplitude of the conductance induced in the postsynaptic cell by a presynaptic spike. In addition, 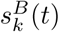 is the synaptic gating variable attached to presynaptic neuron *k* from population *B*, even though it represents changes at the postsynaptic side. It obeys the dynamics described by

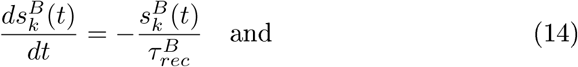

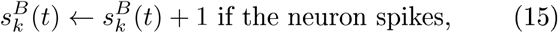

where 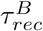 is the synaptic decay time constant. It should be noted that Eq. 13 implies that this quantity is then also the decay time constant of the postsynaptic current. The values for the subtype-specific parameters that were introduced in this paragraph are given in Table 2, which is based on multiple papers that explored subtype-specific properties in mouse V1 [3, 25, 27]. In addition, *σ*^*PC*→*SOM*^ = 1*/*2 and *σ*^*B*→*PC*^ = *σ*^*B*→*PV*^ = 1*/*6 where *B* ∈ {*PC, PV, SOM*}, which reflects the experimental finding that intralaminar projections to PC and PV cells are more spatially restricted than the ones to SOM cells. Furthermore, we set 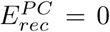 mV and 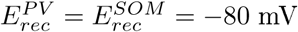. Finally, *S*^*PC*^ = 0.5 and *S* ^*PV*^ and *S*^*SOM*^ were varied to assess their effect on the network’s synchronization. Figure 2A shows a schematical depiction of the network architecture that results from these parameter settings.

**Table 2:** The values for subtype-specific parameters associated with the recurrent inputs. Abbreviations: Presyn. = presynaptic, PV = parvalbumin expressing, Pyr. = pyramidal, SOM = somatostatin expressing.

| Presyn. cell<br>(B) | $P_{con}^{B \rightarrow PC}$ | $P_{con}^{B \rightarrow PV}$ | $P_{con}^{B \rightarrow SOM}$ | $\bar{g}_{rec}^{B \rightarrow PC}$ (nS) | $\bar{g}_{rec}^{B \rightarrow PV}$ (nS) | $\bar{g}_{rec}^{B \rightarrow SOM}$ (nS) | $\tau_{rec}^B$ (ms) |
| --- | --- | --- | --- | --- | --- | --- | --- |
| Pyr. | 0.07 | 0.44 | 0.44 | 0.2 | 0.8 | 0.2 | 2 |
| PV | 1.00 | 1.00 | 0.00 | 2.4 | 2.4 | 0.0 | 5 |
| SOM | 1.00 | 0.86 | 0.00 | 1.6 | 0.8 | 0.0 | 15 |

### Parameter variations

As mentioned before, our model consisted of 3600 PCs, 495 PV cells and 405 SOM cells which were explicitly modelled using Izhikevich’s model of the spiking neuron [17]. They could receive background, visually induced and recurrent inputs. Here, we discuss the different parameter variations that were used to investigate a wide variety of aspects of our model.

First, we wanted to determine which mechanism (ING or PING) generated the oscillations. Therefore, we considered both subnetworks (the pyramidal-PV cell and pyramidal-SOM cell subnetwork) separately, i.e. we set the projection strength of one interneuron sub-type to *S*^*PV/SOM*^ = 0.5 and the other to *S*^*SOM/P V*^ = 0.0. We then varied the drive to the PCs by varying the background spike rate parameter 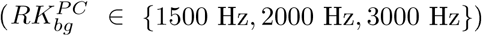 and examined the resulting power density spectra and correlograms. The stimulus size parameter was set to *D*_*vis*_ = 0.0. For all parameter settings, 85 s of network behavior were simulated.

For the following two parameter variations, which were carried out to investigate whether our model could reproduce the attenuation of gamma and the increase of beta power upon visual stimulation (network activation), the projection strengths of the interneurons were set to *S*^*PV*^ = *S*^*SOM*^ = 0.5 and the background spike rate parameter to its standard value 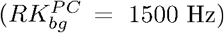. We first varied the stimulus size parameter from 0.0 to 1.0 in steps of 0.1 (*D*_*vis*_ = {0.0, 0.1, …, 1.0}) and examined the results extensively. For each parameter setting, 85 s of network behavior were simulated. We, additionally, wanted to confirm that our model can dynamically reproduce the frequency switching. To do so, we changed the stimulus size parameter during the simulation. Here, the data was acquired in epochs of 5 s. The stimulus size parameter was set to *D*_*vis*_ = 1.0 and we varied the magnitude of the visually induced current. It was set to 0 pA in the first two and the last second, while, during the third and fourth second of the epoch, it was set various values to assess its influence on the network’s synchronization; it could assume values between 20 and 180 pA in steps of 40 pA (*I*_*vis*,0_ = {20 pA, 60 pA, …, 180 pA}). 17 epochs of 5 s were acquired for each parameter setting.

Finally, we wanted to determine for which combinations of PV and SOM cell projection strengths the frequency switching behaviour could be observed. In order to do so, we varied the PV cell projection strength from 0.3 to 0.9 in steps of 0.1 (*S*^*PV*^ = {0.3, 0.4, …, 0.9}) and the SOM cell projection strength from 0.3 to 1.3 also in steps of 0.1 (*S*^*SOM*^ = {0.3, 0.4, …, 1.3}). The network could be either in the resting (*D*_*vis*_ = 0.0) or in an activated (*D*_*vis*_ = 1.0) state. For each parameter setting, 85 seconds of network behavior were simulated. We, furthermore, wanted to investigate in more detail how increasing the PV cell projection strength influenced the network’s dynamical behavior. To do so, the network was simulated for a much longer period during which the PV strength was gradually increased from 0.3 to 0.7 in steps of 0.001 (*S*^*PV*^ = {0.300,0.301, …, 0.7}) while the SOM strength was fixed at *S*^*SOM*^ = 0.5. Again, the network could either be in the resting (*D*_*vis*_ = 0.0) or the activated (*D*_*vis*_ = 1.0) state. 5 s of network behavior were simulated for each setting of the PV cell projection strength and the stimulus size parameter. In all these simulations, the background spike rate parameter was fixed to its standard value 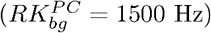.

### Implementation and analysis

The model was implemented using the software package MATLAB R2017a (The Mathworks, Inc., Natick, Massachusetts, United States) and the integration followed Euler’s method. The integration time step size was set to 0.5 ms (sampling rate of *f*_*s*_ = 2000 Hz). The first 5 s of every network simulation were removed prior to analysis so that the initial conditions did not influence the results. Each simulation was repeated for 10 different settings of the random seed, which controlled the random variables in both the realization of network connectivity and the temporal dynamics, to estimate the variance in the results as a consequence of a particular choice of random variables. The mean potentials across the separate neuron subtypes and the spike times were stored for further analysis. We used two measures as LFP estimate 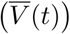 to make sure that observed effect sizes did not critically depend on out choice of LFP model: the mean potential of the PCs and the peristimulus time histogram (PSTH) of the spikes produced by the PCs, which was calculated using a 1 ms bin size (sampling rate of *f*_*s*_ = 1000 Hz). Both of these measures were used in three types of analysis: spectral analysis, the construction of spectrograms and the calculation of spike-LFP pair-wise phase consistencies (PPCs). The PSTHs across the other cell types were also derived from the spike times and were used to calculate cross-correlation functions, so that these functions reflected the relative spike timings of the individual neuron subtypes with respect to one another. The analysis of the recordings was performed using MATLAB’s signal processing toolbox. We will now briefly describe each of the used analysis methods.

For spectral analysis, the spectral power was based on segments with a length of 5 s, which were demeaned by subtracting the mean across time for each individual segment. Subsequently, each demeaned segment was subjected to the multitaper spectral density estimation method with 5 tapers that had a time half bandwidth product of 3 [28]. Finally, the mean was taken across the 16 spectra to obtain the average power spectrum for each setting of the parameters and the random seed.

The calculation of the spike-LFP PPCs required the use of the total length of the recordings; the data was not cut into smaller segments. For the analysis, two regions had to be defined with respect to the square patch. The spikes that were used to determine the PPCs were those corresponding to the neurons from the center area (Figure 2B, black shaded area), while the LFP estimate (mean potentials or PSTH) was calculated on basis of the PCs that were located in a reference region (Figure 2B, orange shaded area). These distinct regions were defined to avoid spurious PPCs. The LFP estimate was demeaned and bandpass-filtered using two filters with distinct frequency bands: one filter only kept the beta band (15 − 25 Hz) and another one only the gamma band (40 − 60 Hz) oscillations. The filtering was performed using an elliptic IIR filter with a passband ripple of 0.1 dB and a stopband attenuation of 60 dB via the MATLAB functions `designfilt` and `filter`. The discrete Hilbert transform was taken of both bandpass-filtered signals [29]. The intermediate result, therefore, comprised two bandpass-filtered analytical signals (signals with both real and complex valued components), one for each frequency band. The phase of all complex values within these two signals were calculated so that estimates for the instantaneous phases (*ϕ*^*β*^ and *ϕ*^*γ*^) were retrieved. The spikes that had been produced by neurons located in the center region were assigned their corresponding phases, i.e. the phases that each of the oscillation types had when these action potentials appeared. Then the spike-LFP PPCs (ϒ^*β*^ and ϒ^*γ*^) were calculated for all neurons separately via [30]

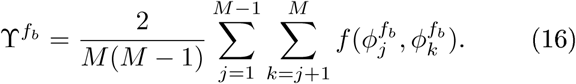

In this expression, 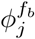 is the phase that was assigned to the *j*^th^ spike from the considered neuron with respect to frequency band *f*_*b*_ ∈ {*β, γ*} and 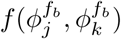 represents the inner product of the two angles:

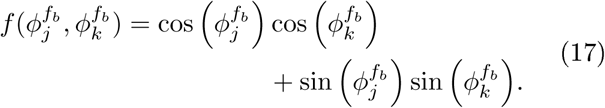

In addition, *M* is the total number of action potentials that were produced by that cell. Finally, the mean was taken across the PPCs from the individual neurons in the center region per subtype to yield the PPC of that group of cells for a given setting of the parameters and the random seed.

Correlation analysis was also performed across the entire length of the recordings, though now the LFP estimates were constructed for each individual cell type by calculating the PSTHs on the basis of all the neurons of that particular type in the network. Each LFP estimate was again demeaned and, from the resulting signal, the cross-correlations between subpopulations (*R*^*B*→*A*^(*τ*_*l*_)) as functions of the lag between the postsynaptic and the presynaptic activity (*τ*_*l*_) were determined via [31, 32]

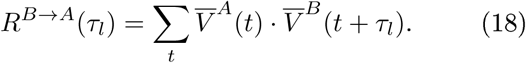

These functions were normalized using the autocorrelations of each separate subpopulation yielding the normalized cross-correlation functions 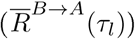:

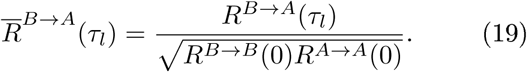

Correlation functions were in some cases subjected to spectral analysis by using the fast Fourier transform (FFT). First the correlation function was transformed and, subsequently, the absolute values were taken of the complex valued outcome. Next, this two-sided spectrum was transformed to a one-sided one. Finally, the values were squared to get the power density spectrum. From such a spectrum, it could be determined to what extent the beta and gamma oscillations were present in the correlation functions.

Spectrograms were constructed to confirm that the frequency changes following visual stimulation could also be obtained dynamically and to determine the range of the PV projection strengths that allowed for this behavior while the SOM cell projection strength was fixed. For the analysis used to confirm the dynamical nature of the frequency switching, a square window function with a size of 200 ms was slid over the time series in steps of 100 ms. As 16 usable 5 s long segments were acquired for each setting of the random seed for this investigation, 49 windows were obtained from each segment. The power density spectrum of each of them was calculated by demeaning the LFP estimates and subsequently subjecting them to the same procedure that was used to calculate the power density spectra of the correlation functions (see above). Finally, the average across these 16 segments was calculated to retrieve the average spectrogram for each setting of the random seed.

The simulations used for the assessment of the PV projection strengths allowing for the frequency switching behavior, yielded 401 usable 5 s long segments per setting of the random seed. Each of these segments reflected the network behavior for a different setting of the PV projection strength. The segments were divided into 5 smaller 1 s long segments. The power density spectra of these smaller segments were calculated by demeaning them and subsequently subjecting them to the same approach that was used to calculate the power density spectra of the correlation functions (see above) and, finally, their mean was taken. Via this procedure, a spectrogram consisting of 401 power density spectra was obtained for each setting of the random seed.

## Results

Our spiking neuron network model comprising 3600 PCs, 495 PV cells and 405 SOM cells is introduced in the Method section. All neurons were spread out across a 2 × 2 square patch of cortex, their membrane potential dynamics were simulated via Izhikevich’s neuron model [17] and each neuron within the network could receive input from three sources at most: the background, the visually induced activity and the recurrent connections. The recurrent connections comprised the inputs the explicitly modelled neurons sent to and received from one another and were initialized according to the scheme depicted in Figure 2A. With regard to the visually induced activity we defined a stimulus size parameter (*D*_*vis*_) which could vary from 0 to 1 and determined the area within the simulated cortical patch in which neurons could receive visual input. This parameter thus approximately reflected the effects of stimulus size, without specifically taking into account any maps present in the mouse visual cortex [21]. By varying the stimulus size parameter as well as the projection strengths of the interneurons, we explored the oscillatory properties of our model. For the specifics of our model and the parameter variations, we refer to the Method section.

In this section, we will discuss the results of this exploration. Firstly, we show that a PING mechanism underlies both the beta and gamma oscillations generated by our model. Subsequently, via the presentation of the results from the stimulus size parameter variation, we demonstrate that our model reproduces the experimentally observed phenomenon of beta power enhancement and gamma power reduction following visual stimulation. Additional simulations confirmed that our model is also able to simulate this frequency switching in a dynamical manner. By means of a grid search with respect to the PV and SOM cell projection strengths, we, however, show that the network only exhibits this frequency switching behavior for a small subset of all possible combinations of these two parameters. Finally, by keeping the SOM cell projection strength fixed and increasing the one of the PV cells in small steps, we investigated the mechanism for switching in more detail.

### Both the pyramidal–PV and pyramidal–SOM cell circuits generate oscillations via a PING mechanism

We first assessed whether the oscillations in the network were generated by means of an ING or a PING mechanism. In order to do so, either the PV or the SOM cells were connected to the PCs and the background activity reaching the PCs was altered. Specifically, we set the strength of one of the subtypes to *S*^*PV/SOM*^ = 0.5 while setting the other to *S*^*SOM/PV*^ = 0.0. The stimulated field parameter was set to *D*_*vis*_ = 0.0 and the background activity to the PCs was varied using 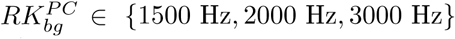. For both circuits, increasing this quantity increased the frequency of the oscillations obtained from the mean potentials of the PCs (Figure 3A–B) but also for those obtained from the PSTHs of the PCs (Figure S1A–B). This increase indicates that the circuits embedded in the model use a PING mechanism to generate the rhythms; if an ING mechanism was involved, the peak frequencies would not have been affected as strongly and hence would have been observed at approximately the same frequencies [20]. The examination of the cross-correlograms of the PSTHs of the PCs and the interneurons confirmed this notion: PCs consistently produced spikes before the interneuron associated with the considered subnetwork (Figure 3C–D), which evidences a PING mechanism underlying the oscillations too [20].

**Figure 3:**
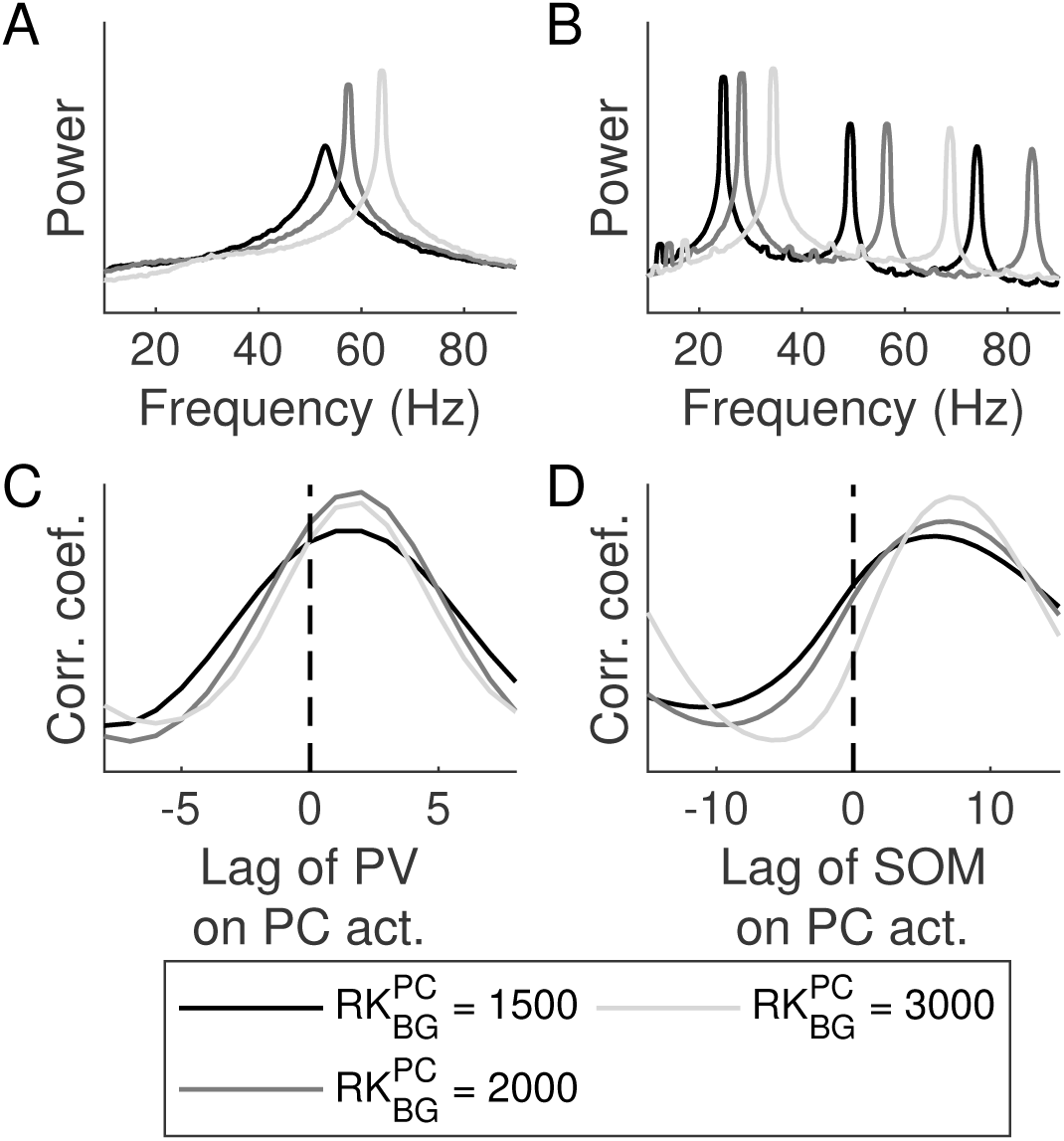
The two circuits comprising pyramidal and parvalbumin (PV) and pyramidal and somatostatin (SOM) expressing neurons, respectively, both generate oscillations via a pyramidal-interneuron gamma (PING) mechanism (A-B) Frequency spectra corresponding to mean potentials of the pyramidal cells (PCs) when considering the pyramidal–PV cell circuit (A) and the pyramidal–SOM cell circuit (B) for various amounts of background input to the pyramidal cells 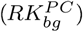. (C–D) Cross-correlogram of the peristimulus time histograms (PSTHs) of the PV and pyramidal cells for the pyramidal–PV cell subnetwork (C) and of the PSTHs of the SOM and pyramidal cells when taking the pyramidal–SOM cell subnetwork into consideration (D). The gray scale represents different amounts of background input to the pyramidal cells 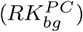. Abbreviations: act. = activation, coef. = coefficient, corr. = correlation

### Visual stimulation enhances beta rhythms via the sculpting of the temporal structure of inhibition

For the next series of simulations we set 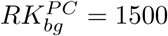 Hz. In addition, the projection strengths of both interneurons were set to *S*^*PV*^ = *S*^*SOM*^ = 0.5. We examined how enlarging the stimulated field, i.e. increasing the value for the *D*_*vis*_ parameter, altered the synchrony in the network activity. The results indicated that the spike rates of the pyramidal and PV cells in the center region (Figure 2B) initially rose but eventually decayed when the extent of the stimulated field was increased (Figure 4A–B). The firing rates of the SOM cells increased monotonously (Figure 4C). These trends are in accordance to the empirically acquired size-tuning curves [4]. Subsequently, we analyzed the power spectra obtained from the mean potentials of the PCs. Three peaks could be observed: a gamma peak at approximately 55 Hz, a beta peak around 18 Hz and another one at roughly 36 Hz (Figure 4D). However, the last of these is the harmonic of the beta peak because it appeared at double that frequency. Considering these spectra for the various stimulated field sizes revealed that the beta and gamma peak were increased and reduced with increasing area of activation, respectively (Figure 4D). This effect was quantified by calculating the average power across the beta (15 − 25 Hz) and the gamma (40 − 60 Hz) frequency bands, which demonstrated that enlarging the area wherein neurons received additional, visual input indeed promoted beta and inhibited gamma oscillations (Figure 4E–F). A consideration of the power spectra obtained from the PSTHs of the PCs confirmed that these findings did not critically depend on the use of the mean potentials of the PCs as the LFP estimate (Figure S2).

**Figure 4:**
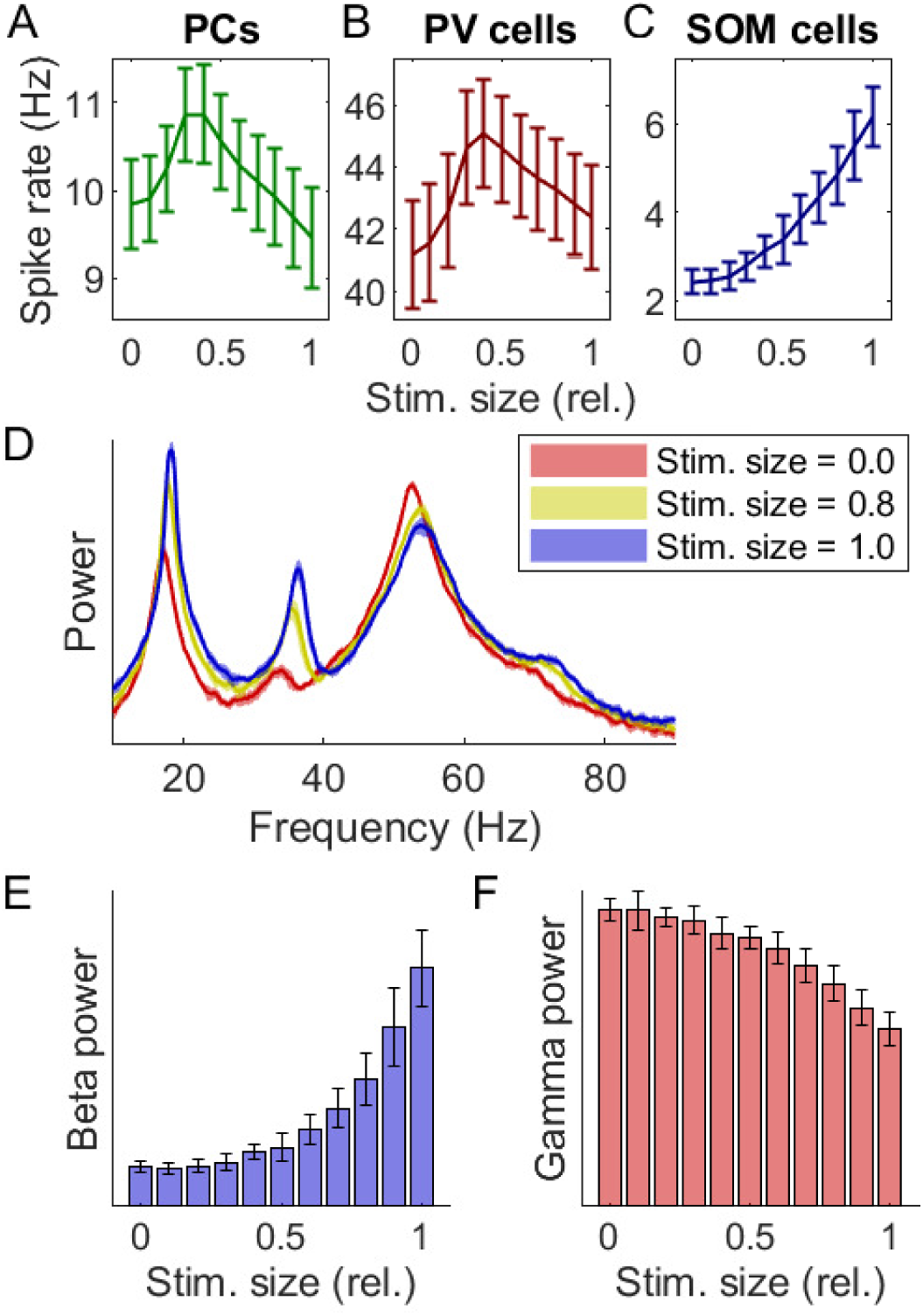
Enlarging the model’s stimulated field elicits a rise and fall of beta and gamma power, respectively. (A–C) Mean spike rates of pyramidal (A), parvalbumin expressing (B) and somatostatin expressing cells (C) in the center region as function of the stimulus size relative to entire simulated area (Figure 2A). Solid lines and error bars depict the *mean* ± *standard deviation* across repeats of the simulation with different random seeds. (D) Frequency spectra for three different relative stimulus sizes. Solid lines and shaded areas depict the *mean*±*standard deviation* across repeats of the simulation with different random seeds. Spectra were obtained by considering the mean potentials of the PCs. (E–F) Mean power across the beta (15–25 Hz, E) and the gamma (40–60 Hz, F) frequency bands as function of the relative stimulus size. Bars and error bars depict the *mean* ± *standard deviation* across repeats of the simulation with different random seeds. The spectra were derived from the mean potentials of the PCs. Abbreviations: rel. = relative, stim. = stimulus.

We wondered how this transition in network synchronization was accomplished. In order to answer this question, the spike time rastergrams were first studied. When none of the neurons received additional, visual input, clear gamma periodic volleys of action potentials originating from the pyramidal and PV cells could be observed (Figure 5A). In contrast, a stimulated field that spanned the whole area elicited strong synchronous, beta rhythmic spiking of SOM cells which caused a strong suppression of pyramidal and PV cells; consequently, the gamma periodic spike volleys produced by the pyramidal and PV cells were less visible (Figure 5B). A larger stimulated field thus appeared to enable SOM cells to produce their spikes more in sync with the beta rhythm. To quantify this, the spike-LFP PPCs of the neurons in the center region were determined for each individual cell type by using the mean potential across PCs as the LFP estimate. The results indicated that all cell types gradually spiked more and less coherent with the beta and the gamma oscillations, respectively, as the stimulated field was increased (Figure 5C–E). The same effect was observed when the PPCs were calculated while using the PSTH of the PCs as the LFP estimate (Figure S3A–C).

**Figure 5:**
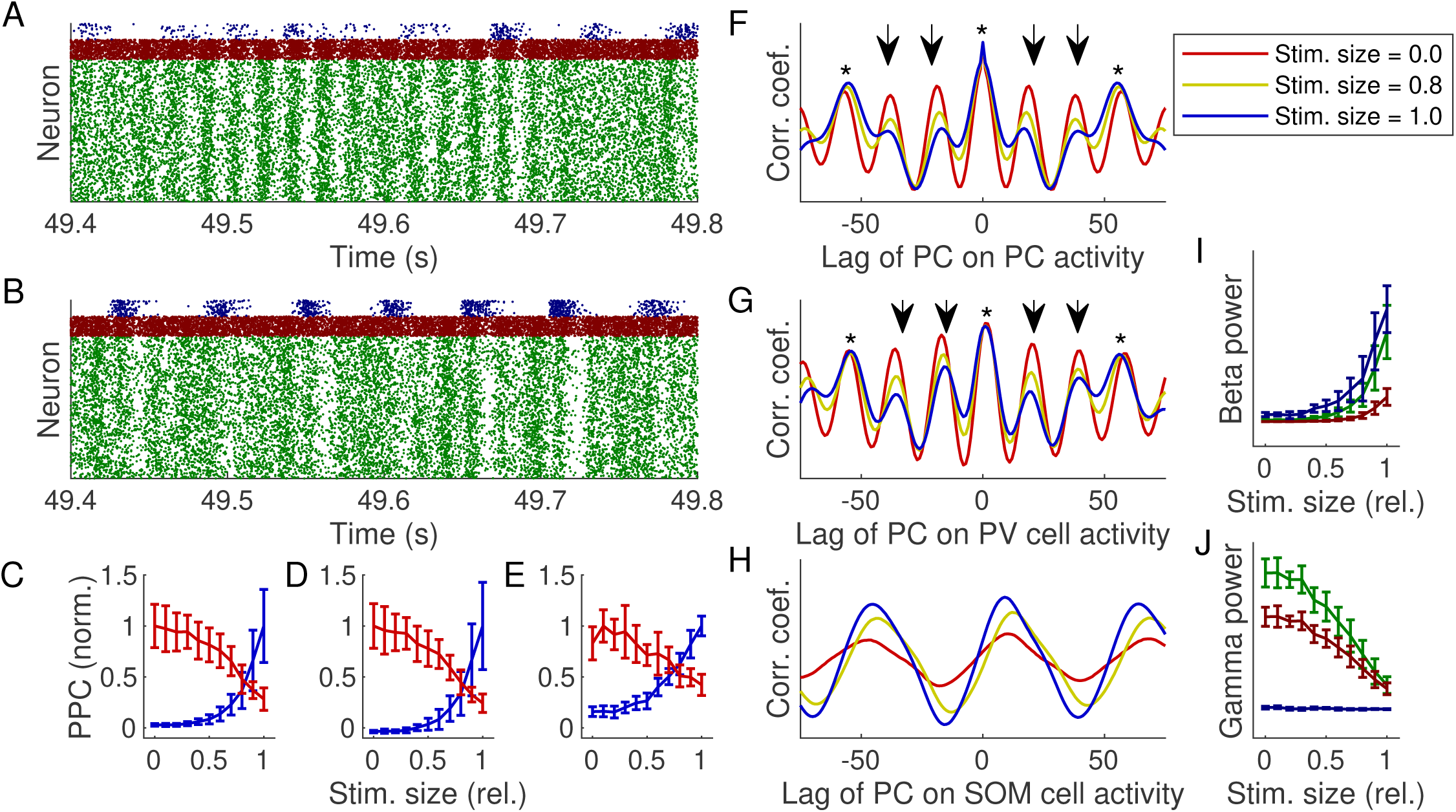
The transition from gamma to beta oscillations for increasing stimulated field size is mediated by the transformation of the gamma oscillation structure by the somatostatin expressing (SOM) interneurons. (A–B) Examples of spike time rastergrams for two stimulated field sizes: *D*_*vis*_ = 0.0 (A) and *D*_*vis*_ = 1.0 (B). Green, red and blue dots represent action potentials from pyramidal, parvalbumin expressing (PV) and somatostatin expressing (SOM) cells, respectively. (C–E) Normalized spike-LFP pair-wise phase consistencies (PPCs) of the pyramidal (C), PV (D) and SOM (E) cells with respect to the beta (15 − 25 Hz) and gamma (40 − 60 Hz) frequency ranges as a function of the stimulus size. Red and blue correspond to the beta and gamma frequency band, respectively. Solid lines and error bars depict the *mean* ± *standard deviation* across repeats of the simulation with different random seeds. PPCs were determined on basis of the mean potentials of the pyramidal cells (PCs). (F–H) Correlograms of the pyramidal cell peristimulus histogram with itself (F) and with those of the PV (G) and the SOM (H) cells for various stimulated field sizes. Common color legend is included in the inset of panel F. Asterisks mark peaks that correspond to beta and gamma rhythms, while downward arrows mark maxima that are only associated with gamma. (I–J) Mean beta (15 − 25 Hz, I) and gamma (40 − 60 Hz, J) power in the different types of correlograms as a function of the stimulated field size. Green, red and blue correspond to the type of correlogram shown in F, G and H, respectively (pyramidal with pyramidal, PV with pyramidal and SOM with pyramidal, respectively). Solid lines and error bars depict the *mean* ± *standard deviation* across repeats of the simulation with different random seeds. Abbreviations: coef. = coefficient, corr. = correlation, pyr. = pyramidal, rel. = relative, stim. = stimulus.

The inspection of the spike time rastergrams and their associated spike-LFP PPCs raised the question as to how the activities of the individual cell types were correlated to one another. Therefore, the correlations between PSTHs of the various cell types in the network were determined. When considering the autocorrelation function of the pyramidal cell activity, it became clear that a larger stimulated field attenuated the extrema in the correlogram that corresponded to a gamma period while the peaks appearing with a beta-like time interval remained relatively unaffected (Figure 5F). The same was observed in the correlogram between the PV cell and pyramidal cell activity, though here the attenuation appeared to be weaker (Figure 5G). The correlation function between the SOM and pyramidal cell activity, on the contrary, seemed to only contain weak gamma periodic peaks, if these were present at all; the corresponding correlogram almost exclusively contained beta rhythms, especially if the stimulated field covered the whole area (Figure 5H). To quantify the presence of both types of oscillations in these correlograms, their power spectra were determined using a FFT. Subsequently, the mean beta (15 − 25 Hz) and gamma (40 − 60 Hz) power were calculated from these spectra for each setting of the stimulated field. The outcome demonstrated that all three correlation functions became more dominated by the beta rhythm as the stimulated field grew, although the PV and pyramidal cell correlogram (Figure 5I, red line) was less susceptible to this change (Figure 5I). A complementary decrease in the gamma power was observed for the correlation functions of the PC activity with itself and with the PV cell activity; in contrast, the correlogram of the SOM with the pyramidal cell activities contained virtually no gamma oscillations for any size of the stimulated field, as was expected (Figure 5J). These results indicate that the SOM cells transform the gamma oscillations that emerge through the interplay between pyramidal and PV cells, so that beta rhythms are amplified at the expense of gamma periodic network activity.

As a final check, we wanted to confirm that our model could also reproduce the frequency switching phenomenon dynamically, i.e. beta and gamma power could be enhanced and reduced, respectively, through activation of the network within the same simulation time series. In order to do so, we set the activated area to cover the entire simulated cortical patch (*D*_*vis*_ = 1) and altered the visually induced current (*I*_*vis*,0_) within a 5 s epoch: this parameter was set to 0 pA in the first two seconds and the last second and was set to varying values in the intermediate two seconds. Inspection of the spectrograms constructed on the basis of the mean potentials of the PCs revealed that beta and gamma power were indeed in- and decreased, respectively, when the visual current was higher than zero (Figure 6A–E). The variation of the magnitude of the current associated with the visual stimulation, furthermore, demonstrated that higher values for this parameter yielded larger power differences following visual stimulation (Figure 6A–E).

**Figure 6:**
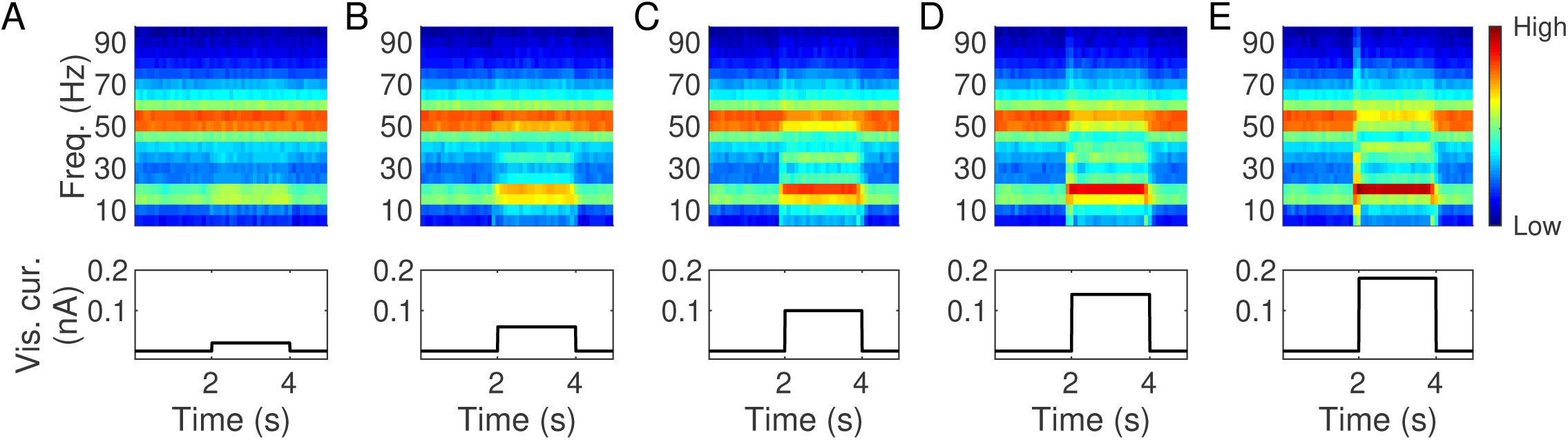
The frequency switching behavior can be induced dynamically in the proposed spiking neuron network model. (A–E) Spectrograms of the mean potentials of the PCs (top row) and the concurrent change in the visual current (*I*_*vis*,0_) throughout an epoch (bottom row). The latter plots show that this parameter was 0 pA in the first two and the final second of the epoch and could have a variety of values in the third and fourth second, namely 20 pA (A), 60 pA (B), 100 pA (C), 140 pA (D) or 180 pA (E). In the spectrograms, black and white colors correspond to a low and high power, respectively. The same color scale has been used for all spectrograms in this figure.

These results, which were also observed for the spectrograms acquired from the PSTHs of the PCs (Figure S4), thus demonstrated that our model could indeed reproduce the frequency switching phenomenon in a dynamical manner and additionally indicated that, in our model, the extent to which the beta and gamma power are amplified and attenuated, respectively, not only depends on the stimulus size parameter, but also on the magnitude of the network activation current (*I*_*vis*, 0_).

### In order to amplify beta and attenuate gamma oscillations by means of visual stimulation, the PV and SOM cells must be allowed to compete for oscillatory control of the PCs

We wanted to find out how generic the network motif was that allowed beta oscillations to outcompete gamma upon visual stimulation. We, therefore, studied the conditions under which the dominant beta and gamma oscillatory activity via the enlargement or shrinkage of the spatial extent of the visual stimulation occurred by varying the projection strengths of the PV (*S*^*PV*^) and the SOM (*S*^*SOM*^) cells while either none of the neurons received input through visually induced activity (the rest state network) or the stimulated field size covered the entire square patch. At first, we looked at the mean spike rates of the individual cell types. In addition to the pyramidal and PV cells being suppressed by stronger PV and SOM cell projections, their firing rates also exhibited a local minimum (Figure 7A–B). The SOM cells, on the other hand, show a very different pattern: higher levels of SOM cell inhibition increased their firing rate while stronger PV cell projections reduced it (Figure 7C). The beta power follows the same pattern as the SOM cell firing rate (Figure 7C–D), while the trend of the gamma power has more in common with those of the pyramidal and PV cells (Figure 7A–B,E). These spectral densities were determined for the rest state network on the basis of the mean potentials of the PCs. Also the power that corresponded to a stimulated field that covered the entire square patch was determined using that same measure as the LFP estimate and, subsequently, the difference between the power associated with the visually activated and resting state network was calculated. We observed that there is only a small subset of settings for the PV and SOM cell projection strengths that gives rise to higher beta and lower gamma power in the activated state (Figure 7F–G). Similar findings were obtained when the PSTHs of the PCs were used as the LFP estimate (Figure S5). We determined which settings brought about each kind of behavior. The results of this assessment demonstrated that there is only a limited region with regard to the PV and SOM cell projection strengths that gives rise to enhanced beta and decreased gamma oscillations via activation of the network (Figure 7H). Gamma and beta rhythms dominated the network’s oscillatory activity when the strength of the PV and SOM cell efferents, respectively, were too strong (Figure 7D,E,F,G,H).

**Figure 7:**
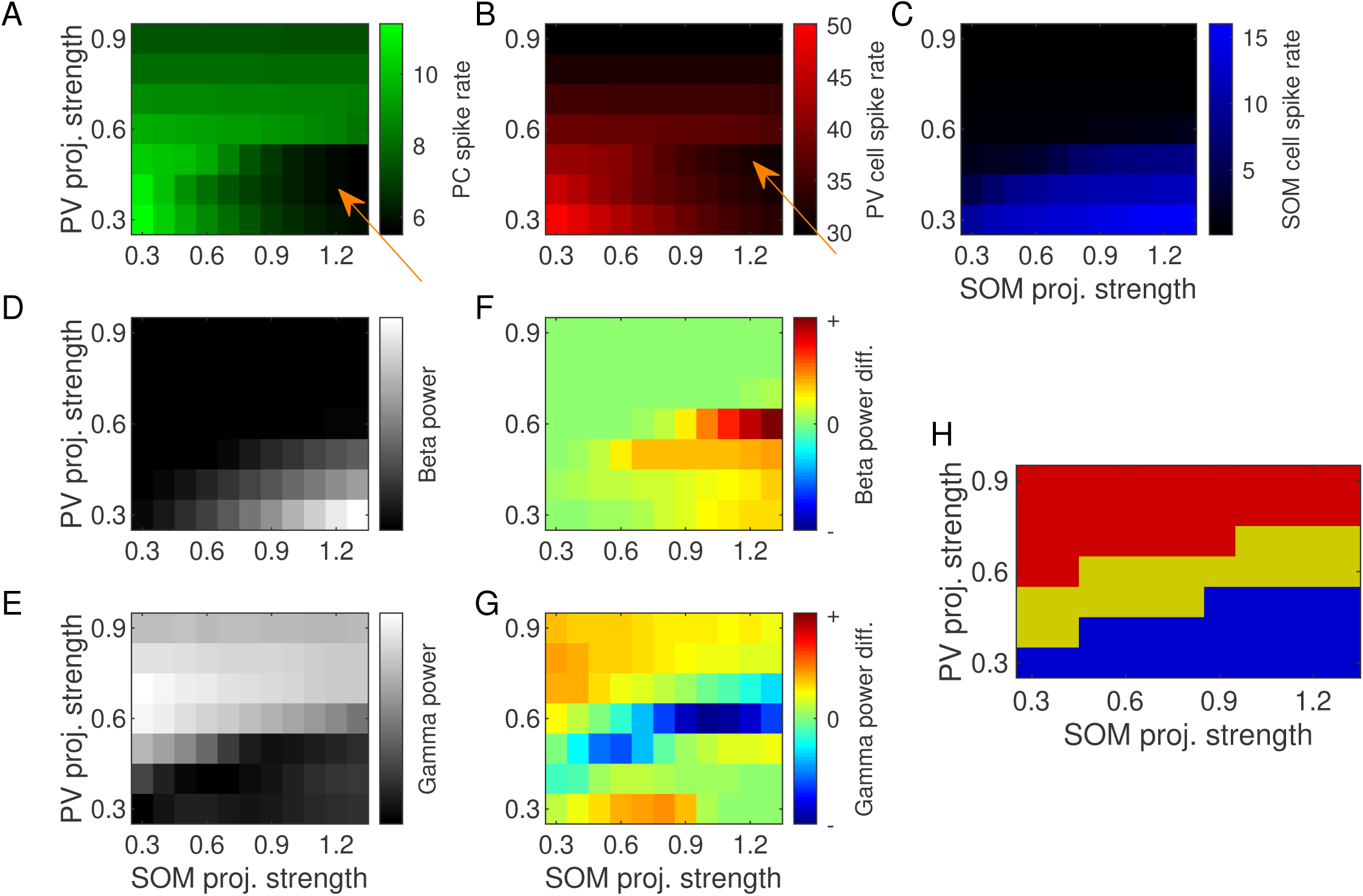
Visual stimulation can induce an enhancement of beta and a reduction of gamma power for a restricted range of parvalbumin expressing (PV) and somatostatin expressing (SOM) interneuron projection strengths. (A–C) Mean firing rates of pyramidal (A), PV (B) and SOM (C) cells in the center region as a function of the projection strengths of the two interneuron types. Orange arrows point to local minima. No neurons received additional input through visually induced activity for these panels. (D–E) Beta (D) and gamma (E) power as a function of the PV and SOM cell projection strengths. For these plots, none of the neurons received additional input through visually induced activity. The frequency spectra were obtained from the mean potentials of the PCs. (F–G) Difference in beta (F) and gamma (G) power between the situation wherein the stimulated field covered the entire square patch and the one without any additional visual input. A positive difference means that the power is larger for visual stimulation. Powers were derived from the frequency spectra obtained from the mean potentials of the PCs. (H) Schematic depiction of the projection strength settings that resulted in the different types of network synchronization, i.e. network synchronization behavior as a function of the PV and SOM cell projection strengths. Red and blue colors correspond to the network persistently oscillating in the gamma and beta frequency band, respectively, while yellow represents a system being able to switch between these two types of rhythms via activation of the network. Abbreviations: diff. = difference, proj. = projection.

These findings indicated that the system requires some sort of competition between PV and SOM cells over the oscillatory control of the PCs in order for it to alter between synchronization in the beta and gamma frequency bands via activation of the network. To support this hypothesis, we performed additional analyses. First, the spike time rastergrams were examined. These showed that when the gamma oscillations were persistent, the number of SOM cell action potentials was very low, and as a consequence, the PCs spiked periodically with a typical gamma period, regardless of the amount of activation (Figure 8A–B). However, the PV cells still spiked when persistent beta oscillations were observed, but in that case PCs nevertheless produced action potentials synchronized to a beta rhythm (Figure 8C–D). When network activation can induce enhanced beta and decreased gamma rhythms, the additional input to the PCs apparently leads to suffcient collateral depolarization of the SOM cells and, consequently, the PCs’ periodic spiking is altered (Figure 8E–F). We quantified the different types of PC periodic firing by means of the spike-LFP PPCs calculated using the mean potentials of the PCs, which revealed the behavior inferred from the spike time rastergrams: beta and gamma rhythm associated PPCs were in- and decreased, respectively, following network activation, especially when the parameter settings were considered that gave rise to the shifts in main network oscillation frequency upon an increased spatial extent of the visual stimulation (Figure 8G–J). Similar findings were observed when the spike-LFP PPCs were calculated on the basis of the PSTHs of the PCs (Figure S6).

**Figure 8:**
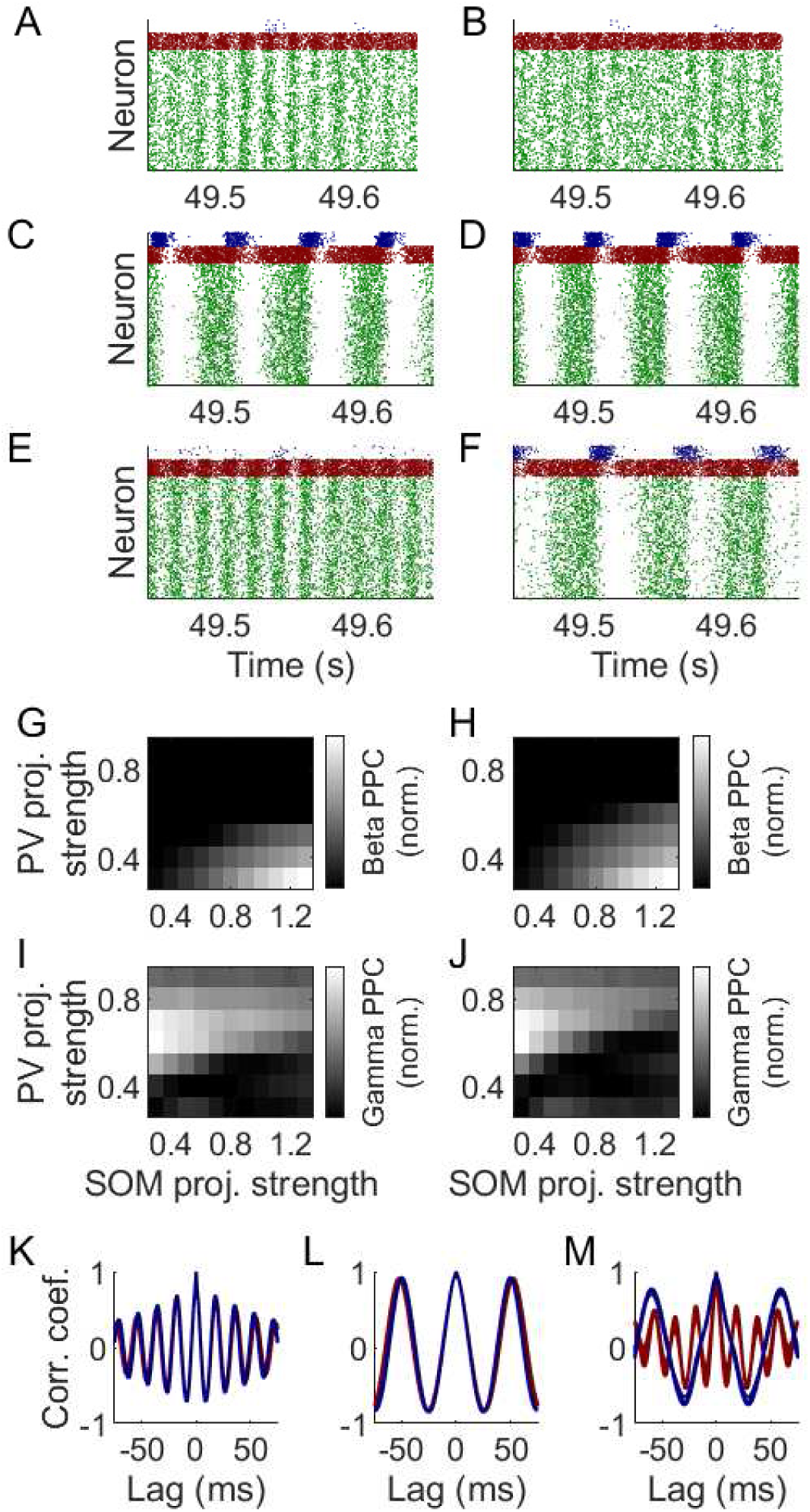
Higher beta and lower gamma power induced via network activation requires a balance between the inhibitory contributions of parvalbumin expressing (PV) and somatostatin expressing (SOM) cells and their associated oscillatory control over the pyramidal cells (PCs). (A–F) Spike time rastergrams for the rest state (left column) and the activated network (right column). Green, red and blue dots represent action potentials of pyramidal, PV and SOM cells, respectively. For these plots, the PV and SOM cell projection strength were set to 0.9 and 0.3 (A–B), 0.3 and 0.9 (C–D) and 0.6 and 1.2 (E–F). (G–J) Spike-LFP pair-wise phase consistencies (PPCs) of the PCs for the rest state (left column) and the activated network (right column) with respect to the beta (G–H) and gamma (I–J) frequency bands. PPCs were obtained using the mean potentials of the PCs as LFP estimate. (K–M) Correlograms for the PC to PC connections when the PV and SOM cell projection strength were set to 0.9 and 0.3 (K), 0.3 and 0.9 (L) and 0.6 and 1.2 (M). Red and blue correspond to the rest state and activated network, respectively. Abbreviations: coef. = coefficient, corr. = correlation, norm. = normalized, proj. = projection.

Finally, we examined the correlograms of the PC to PC connections, since it was shown earlier that network activation attenuated the gamma rhythm associated secondary peaks in these graphs. If gamma or beta oscillations dominated the response, i.e. if these rhythms were persistent across network activation conditions, no substantial differences could be observed in the auto-correlations of the PCs (Figure 8K–L). On the other hand, when a main oscillation frequency shift was inducable, the correlogram should be drastically transformed; gamma rhythm associated peaks were almost completely repressed when the network was activated, while the extrema corresponding to a beta oscillation were amplified by such an event (Figure 8M).

When considering Figure 7 and Figure 8 in the context of the emergence of the spontaneous gamma oscillations during the CP, it must be recognized that the juvenile mouse V1 is represented by a network configuration similar to the blue region in Figure 7H: the LFP of the juvenile mouse V1 also mostly contains beta oscillations in both the resting as well as the activated state. The V1 of adult mice, on the other hand, has network dynamics resembling the yellow area in Figure 7H, because the mature spontaneous LFP primarily contains gamma oscillations while beta oscillations are predominantly found in its visually evoked counterpart. Our results therefore imply that the influence of PV cells within the network increases across the critical period of mouse V1. Given this observation, we wanted to obtain a more detailed overview of how the PV cell projection strength influences the network dynamics when the SOM cell projection strength was fixed.

For this investigation, we set the SOM cell projection strength to *S*^*SOM*^ = 0.5. The PV cell projection strength was varied from 0.300 to 0.700 in steps of 0.001 (*S*^*PV*^ ∈ {0.300, 0.301, …, 0.700}). However, we changed the parameter during one simulation time series. More specifically, it was initially set to 0.300 and was increased with the mentioned step size after every 5 s of simulation until the upper limit of 0.700 was reached. We determined the power density spectrum using the mean potentials of the PCs for each of these epochs and plotted them as a function of the PV cell projection strength. As expected, the results demonstrated that the power in the beta and gamma frequency band were reduced and enhanced, respectively, for increasing PV cell projection strengths, both when considering the resting-state and activated network (Figure 9A–B). However, they also showed that a distinct range of PV cell projection strengths existed wherein beta and gamma powers were in- and decreased following network activation, respectively (Figure 9A–B). To identify this range in more detail, we calculated the mean beta (15–25 Hz) and gamma (40–60 Hz) power for each setting of the PV cell projection strength and, subsequently, determined the difference between the powers corresponding to the activated and the resting-state network. The resulting plot clearly indicated that the beta power was never lower for the activated situation than for the resting-state (Figure 9C). The gamma power, on the other hand, could be lower, though only for a restricted range of PV cell projection strengths (Figure 9C). One may argue that, as this parameter increases from 0.300 to 0.700, the system goes through two ‘bifurcations’: one around 0.410 and another around 0.590 (Figure 9C). At the former, the system’s dynamics transition from a beta oscillation dominated system to a balanced state with regard to the inhibitory contributions of both interneuron types wherein the frequency switching phenomenon can be induced; at the latter, the network dynamics exit this balanced region and enter a regime in which gamma oscillations govern both the spontaneous as well as the activated state. Similar results were found when the PSTHs of the PCs were used as the LFP estimate (Figure S7). Finally note that these findings not only provide a more explicit investigation of the system’s dynamical behavior as a function of the PV relative to the SOM cell projection strength, but also demonstrate that an increase of the influence of the PV cells in the network over time may indeed underlie the emergence of spontaneous gamma oscillations during the CP; after all, any network synchronization changes induced via the augmentation of the projection strength of these interneurons were obtained dynamically as this parameter was incremented within one and the same simulation time series of the network’s dynamics.

**Figure 9:**
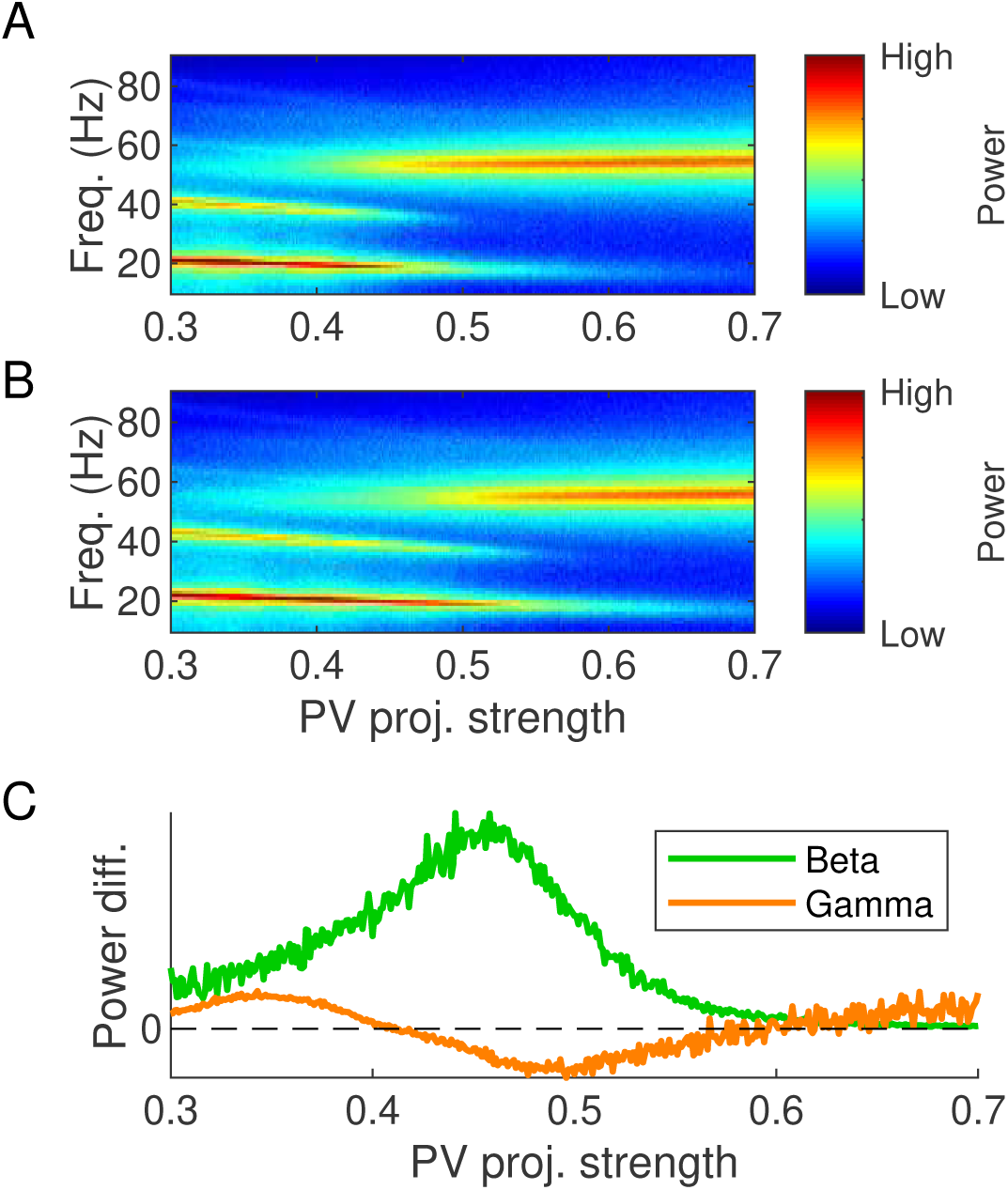
A detailed investigation of the network dynamics for varying parvalbumin expressing (PV) cell projection strengths provides an overview of the ‘bifurcation’. (A–B) Power density spectra of the resting-state (*D*_*vis*_ = 0, A) and the activated (*D*_*vis*_ = 1, B) network as a function of the PV cell projection strength for a fixed somatostatin expressing (SOM) cell projection strength (*S*^*SOM*^ = 0.5). The mean potentials of the PCs were used for the derivation of the spectra. (C) Power differences between the activated and resting state network activity with respect to the beta (15–25 Hz, green) and gamma (40–60 Hz, orange) frequency band as a function of the PV cell projection strength for the same fixed SOM cell projection strength. A positive and negative difference reflect a higher and lower power in the activated situation, respectively. Powers were calculated on the basis of the mean potentials of the PCs. Abbreviations: diff. = difference, freq. = frequency, proj. = projection.

## Discussion

In summary, our model, which generates beta and gamma oscillations via a PING mechanism (Figure 3), is able to reproduce the empirically-observed switch from gamma-to beta-dominated synchronization following visual stimulation (Figure 4). This shift is caused by the SOM cells transforming the gamma oscillations so that the network generates beta rhythms instead (Figure 5). Finally, we have shown that this can only occur when PV and SOM cells are allowed to compete over the oscillatory control of the PCs (Figure 7–9). These results still require an interpretation in the context of the model that has been introduced in this article and the available experimental literature. In the following, we discuss the relevance of this study and any future prospects that will arise from it.

### PING is a plausible mechanism for oscillation generation and observed frequency shifts in the V1 of mice

In our model, the SOM cell associated beta and PV cell associated gamma oscillations are both generated via a PING mechanism (Figure 3A–D). However, the translation of this result to the experimental context is not immediate. Although most of the parameters have been directly derived from the experimental literature, those corresponding to the background currents are more difficult to infer. The background current to a neuron can be regarded as the summation of the inputs coming from more distant neurons and overall level of neuromodulators. This mix of factors makes it rather difficult to assign meaningful values to the parameters governing this quantity.

Nevertheless, the results have been included here to show how our model generates oscillations. The resemblance that the remainder of the results has with the experimental data additionally supports the notion that both types of rhythms indeed are being produced by a PING mechanism. We have not evaluated whether ING mechanisms could produce the same synchronization behavior that has been observed in this study and experimental research. Hence, additional research is required to unambiguously identify the underlying oscillation mechanism.

### The relation between the stimulated field size and the induced rise in beta and fall in gamma power

Our model reproduces the experimentally observed enhancement of beta and reduction of gamma power in the LFP of mouse V1 following the presentation of a visual stimulus to the animal (Figure 4E–F) [5, 6]. Additionally, the simulations also reproduce the subtype-specific size tuning curves described by the literature (Figure 4A–C) [4]. However, one must be careful in interpreting these findings. In the model, the stimulated field size is a relative measure that cannot be related directly to a physical stimulus size. The results presented in this article, therefore, demonstrate that, when the magnitude of the visually induced current is fixed (Figure 6A–E), the activated area of V1 determines to which extent beta oscillations are induced in the visually evoked LFP. The model thus imitates electrophysiological properties of mouse V1 through its biologically plausible connectivity patterns, which have been derived from the experimental literature.

Additionally, our results show that the enhancement of the beta rhythms at the expense of the gamma oscillations is accomplished via outcompeting the PV-PC subnetwork by the SOM-PC subnetwork (Figure 5F–J). The findings, furthermore, imply that the main beta peak appears at approximately a third of the gamma peak frequency (Figure 4D), which is consistent with experimental data as well [5, 6, 9]. We have also shown that too strong PV and SOM cell projections result in persistent gamma and beta oscillations, respectively (Figure 7H,Figure 9C). Hence, the switch in main synchronization frequency upon visual stimulation requires a competition between PV and SOM cells and their oscillatory control over the PCs (Figure 8A–F). Competition also reduces the spike rates in the network (Figure 7A–C). When the PV projections are too powerful, SOM cells are almost completely silenced, whereas the PV cells still remain active when the SOM cell projections are too strong. Our results therefore indicate that the visually evoked emergence of beta oscillations is a consequence of the PV cells being unable to effectively suppress the PCs. This enables the PCs to activate the SOM cells which, in their own turn, impose the network to oscillate in the beta frequency range.

The function of this switch in main oscillation frequency is not fully understood, but some ideas have been presented. For instance, it has been argued that beta oscillations in primates are related to the maintenance of the current cognitive state [33]. More interestingly, it has been shown in rodent that, because of their horizontally aligned afferents [4], SOM cells promote synchronization across cortical space [6, 10]. This property of the SOM cells and the fact that in our model SOM cells are activated when PV cells are unable to effectively suppress PC activity together imply that strong visual stimulation triggers the generation of beta oscillations so that also more distant cortical areas become increasingly synchronized with the V1. The latter, in its own right, would then putatively improve the information transfer between cortical areas. Though improved information transmission is typically observed for gamma oscillations [34], the beta oscillations considered in this paper may facilitate this as well for their peak frequencies are intermediate between the experimentally determined beta and gamma frequency bands.

### Network configuration alterations may explain the emergence of gamma oscillations during the critical period

The motivation for this study was to find the configurations of a spiking neuron network model that are responsible for V1 dynamics in juvenile and adult mice. Specifically, we aimed to find the connectivity changes explaining the establishment of spontaneous, high-frequency gamma oscillations, which are disrupted at the benefit of the beta rhythms following visual stimulation, in mouse V1 during the CP [5]. Since stronger PV cell projections were found to increase the gamma power (Figure 7E), our results indicate that an overall strengthening of PV cell projections across this time window may very well be the reason for the emergence of these rhythms. At the same time, the outcomes of our simulations and analyses also demonstrate that the PV and SOM cell associated influences on the PCs should be balanced at the end of the CP. It is exactly after this period of enhanced plasticity that gamma power should be suppressed and beta power augmented during visual stimulation of the mouse [5] and our model only exhibits such behaviors for a restricted range of PV and SOM cell projection strengths (Figure 7H). Therefore, this study shows that, during the CP, PV cell inhibitory contributions become stronger until the network reaches that balanced state.

Plasticity mechanisms are one method to reinforce these projections and additional experimental evidence supports the notion that PV cell related plasticity underlies the enhancement of gamma powers during the CP. For instance, the opening of the CP has been linked to the maturation of a subset of the GABAergic interneuron population [8, 35] and there is evidence that that maturing subset comprises the PV cells. When stem cells derived from the medial ganglionic eminence, the embryonic brain region that produces PV and SOM cells during development [36], are transplanted into the V1 long after the CP, they differentiate to a large extent towards this interneuron subtype and functionally integrate themselves into the host network [11, 12]. A consequence of this transplantation and subsequent integration is the putative induction of a time window with enhanced plasticity that resembles the CP [11]. Likewise, the closure of the CP is marked by molecular and cellular advancements too. The appearance of molecular ‘brakes on plasticity’, like myelin sheaths, that have Nogo-A as an associated protein, and the PNNs, that were mentioned in the Introduction, namely coincides with the end of the CP [13, 37, 38]. Especially the latter type of consolidators, the PNNs, has recently gained much interest in multiple studies [7, 15, 13, 39]. It has been shown that these nets primarily enwrap PV cells and that their removal reactivates ocular dominance plasticity in the V1 of mice [13]. Additionally, more recent studies have demonstrated that PNN removal also increases gamma power right after, but not for longer periods of, MD [7], that it disrupts the retrieval of remote fear memory [39] and that in L4 it leads to increased thalamic PV cell recruitment [15].

Nevertheless, other explanations for the increase of the gamma oscillations during the CP are possible. In other brain areas, it is, for example, known that the decay time constant of the IPSC of PV cells declines during development [40, 41]. Since one of these studies investigated the barrel cortex of mice, which shares many developmental aspects with the V1 [42], this potential mechanism should be assessed. However, a quick, mathematical evaluation of the effect that such a development would elicit, reveals that it only further weakens the influence that PV cells have on the PCs. Another, more promising study has found that the SOM cells lose cholinergic responsiveness during the CP, which would lower their excitability [43]. As a consequence, these cells would have weaker control of the PCs and, complementarily, the influence of the PV cells on the excitatory cells would increase. This developmental loss may therefore evidently contribute to the emergence of gamma oscillations during the CP.

In summary, the CP thus seems to be marked by high amounts of PV cell related plasticity and our results provide a new insight as to how this plasticity may change the network. Experimental data, for example, suggest that PV cell projection strengthening already occurs right before the onset of the CP and a weakening is observed with respect to the SOM cells [44], but what exactly happens with these quantities during the CP could not be determined. Here we have provided support for the idea that PV cell projections are strengthened and that the cells themselves become integrated in the circuit of the V1 during this time window. Secondly, this study indicates that the plasticity mechanisms that are at play during the CP aim to eventually find a network configuration that results in PV and SOM cells competing over the oscillatory control of the PCs. We even have shown that in our model this development can be replicated by increasing the PV cell projection strengths over time (Figure 9A–C). These findings could be exploited in future studies to devise therapies that reverse the effects of early onset inherited retinal dystrophy.

### The precise function of the emergence of gamma oscillations during the CP remains unknown and its unraveling requires more study

The question still remains as to what is the function of the emergence of the gamma oscillations during the CP. Is their appearance and subsequent fading over the next 24 hours following MD during the CP or after PNN removal [7] just an epiphenomenon or do they fulfill a particular function? A related experimental study at least provides evidence that this finding is in line with our outcomes. In juvenile mice, PC activity drops right after the mouse is monocularly deprived, but returns to its original level in the 24 hours that follow; this return is facilitated by a decrease in PV cell activity [14]. The link between these two experimental studies and our results should by now be clear: all three demonstrate that relatively large PV cell activity enhances gamma power and that restoration of the PC activity level via PV cell projection weakening restores the desired balance within the inhibitory contributions from the different interneurons.

In addition, gamma rhythms have been found to induce plasticity mechanisms [45]. In humans, they are, for example, associated with working and short-term memory [46]. Studies of non-human primates have, furthermore, shown that a stimulation pulse train that is phase-locked to a beta oscillation induces short-term potentiation and depression of the connections if the pulse train is phase-locked to the depolarization and hyperpolarization phase of that rhythm, respectively [47]. The strengthening or weakening is assumed to be facilitated by a spike-timing dependent plasticity (STDP) mechanism [47], which is a type of Hebbian learning that considers the relative timings of the spikes of two neurons in order to decide whether their connection should be stronger or weaker [48, 49, 50]. Oscillations may thus regulate the timing of spikes so that the synaptic transmission efficiency between two neurons is appropriately adjusted via STDP.

However, to our knowledge, oscillations have, so far, not been linked to CP plasticity. The current consensus is that its mechanisms affect the inputs from both eyes independently during MD [51, 52]. Specifically, the inputs from the deprived eye are weakened in the first two days of MD, most likely via long-term depression, while those corresponding to the open eye are amplified via homeostatic pathways in the subsequent five days [53, 54, 55]. More experimental and theoretical assessments are needed to determine whether any of these distinct plasticity mechanisms critically depend on the equilibrium between PV and SOM cell inhibitory contributions, which we have shown is required for proper V1 dynamics after the CP. If so, additional research should be devoted to discover whether the emergence of gamma oscillations during the CP is simply an epiphenomenon or whether these rhythms actually play a role in CP plasticity.

### How our model relates to other computational studies to mouse V1

In the Introduction, some neural mass models of mouse V1 were already mentioned. These models successfully reproduced the phenomenon of surround inhibition and the increased beta and attenuated gamma power upon visual stimulation [6, 16]. Though firing rate models may be used to study neural oscillations, it must be acknowledged that spike timing is a determining factor in the generation of LFP signals. Moreover, it has been demonstrated that firing rate and synchrony can be modulated independently, which makes neural mass models less fit to study oscillations [56].

By using a spiking neuron model, we have obtained new insights regarding the beta and gamma rhythms and the roles that PV and SOM cells play in them; specifically, our results indicate that the relative PV and SOM cell inhibitions should satisfy a particular constraint at the end of the CP for a proper functioning of mouse V1. More generally, we have shown that the main synchronization frequency of oscillations generated via a PING mechanism can be altered via network activation. It has already been demonstrated that such an effect cannot be observed when the network hosts a combination of ING and PING mechanisms: in that situation, the highest frequency of the two then dominates the synchronization [57]. Whether ING mechanisms can facilitate peak frequency shifts upon network activation is unclear. Here, it must also be mentioned that the extent to which PCs are involved in the generation of oscillations in the neocortex should be limited in terms of the number of spikes per cycle; as a consequence, it is believed that neural rhythms in the visual system are produced neither purely by a PING, nor purely by an ING mechanism [58]. Note that this does not rule out the applicability of our study to mouse V1: it merely places it in a more nuanced perspective.

To our knowledge, this is the first study of oscillations in V1 during the critical period that involves the explicit modeling of three distinct neuron types. Spiking neuron models that were inspired by this cortical region investigated other properties. One of these, for example, proposed a possible mechanism as to how orientation selectivity can be established in cortices that lack an organized map with regard to this feature [19]. This model merely comprised two neuron classes: excitatory and inhibitory neurons. Another example investigated how stimulus detection performance can be enhanced in noisy spiking neural networks; this model consisted of the same neuron types as have been included in this study [59]. Still, we are not the first to investigate the coexistence of oscillations through a spiking neuron network model comprising three distinct cell types: one model that was based on the hippocampus already demonstrated that coexistence of *θ* and gamma oscillations requires a balance in the effective strengths of the different inhibitory neurons in the network [60]. Our work shows that the same principle is applicable to the beta and gamma rhythms in mouse V1 and additionally demonstrates that network activation can alter the synchronization of the neural ensemble too.

There are multiple types of interneurons, of which the ones classified as parvalbumin positive (PV), somatostatin positive (SOM) and vasoactive intestinal peptide positive (VIP) have received most attention [1] (note that there are alternative labels in use for each of these types). Optogenetic approaches to transgenic animals in which specific cells are either labeled by GFP or express Cre have elucidated the functional role of each type and identified structural motifs in different cortical layers, see for a perspective [61], [62] or [63]. These motifs need to be developmentally established, and this may happen both within critical periods as well as outside. The vagueness of this description derives from the fact that the development of these motifs has not been studied extensively. Here we interpret our simulation results in terms of motifs and the ocular dominance critical period experiments that have been reported in the literature.

Even though there are many types of interneurons, we focus on two groups: the PV and the SOM cells. The literature on CP plasticity identify PV neurons as prime actors [8], and electrophysiological literature identify SOM cells as prime actors in visually induced beta oscillations and horizontal projections mediating surround inhibition [6, 9]. This means we omit from the model VIP interneurons, which do, however, play an important role in the effects of locomotion on visual responses [64] and have a specific neuromodulator sensitivity [65] while defects in their function influence other types of cortical plasticity relevant for cognitive function [65]. We will defer a computational investigation of their role in the context of modulating oscillations to a future study. Here, the model in [66] may serve as inspiration.

## Conclusion

In this article, the differential roles of PV and SOM cells in the generation of oscillations have been investigated. From our results, three main conclusions can be drawn. First of all, the emergence of gamma oscillations during the CP [5] is most likely caused by an overall increase in the influence that PV cells have on the PCs in the network. Given the currently available knowledge of the CP, plasticity presumably underlies this development, which would concretely imply a general strengthening of PV cell projections across this time window. Secondly, this increase in influence has a limit: persistent gamma oscillations emerge if PV cells become relatively too powerful. This would prevent visually stimulating the animal from inducing the SOM cell associated beta rhythms in the V1, which, as the available literature demonstrates, should actually be possible [5, 6]. Hence, the inhibitory contributions of PV and SOM cells must be balanced at the end of the CP in order for spontaneous gamma and visually evoked beta oscillations to coexist in the V1. Finally, we have presented evidence for a mechanism by which these visually evoked beta oscillations are realized. The results of this study namely indicate that SOM cells transform the dynamic circuit motif laid out by pyramidal and PV cells for the production of gamma oscillations so that it then produces beta oscillations instead. In addition, it has been argued that this implies that beta rhythms emerge when the PV cells are unable to effectively suppress the PCs before they collaterally activate the SOM cells.

In conclusion, our study links many experimental studies together into one comprehensive model that has biologically plausible connectivity patterns. It also provides new insights into how specific members of neural ensembles in the brain can be mobilized to produce different types of oscillations. Furthermore, it demonstrates that experimental observations in electrophysiological studies may be explained by mechanisms that are sensitive to a precise parameter setting and presumably require careful fine-tuning of the network configuration in order to emerge and be maintained.

## Acknowledegments

We thank C. Bollen and M. J. ter Wal for their comments and suggestions on the manuscript. This study fell under the project: ‘Light after dark; restoring visual perception in inherited retinal dystrophies’ (NWO 058-14-002).

