## Supplementary Method and Figures for "Balance between inhibitory cell types is necessary for flexible frequency switching in adult mouse visual cortex"

In [17], Eqs. 2-4 are presented in the following form

$$\frac{dv}{dt'} = 0.04 \cdot v^2 + 5 \cdot v + 140 - u' + I', \tag{S.1}$$

$$\frac{du'}{dt'} = a' \cdot (b' \cdot v - u'), \tag{S.2}$$

$$\text{if } v \geq 30 \text{ mV, then } \begin{cases} v \leftarrow c' \\ u' \leftarrow u' + d' \end{cases} . \tag{S.3}$$

Although it is mentioned in [17] that $v$ and $u'$ are dimensionless variables, the author uses the mV notation when it comes to the variable representing the membrane potential and also explains that $v$ and $t'$ have mV and ms scales. We, therefore decided to consider this spiking neuron model from a physically meaningful, i.e. dimensional, perspective. However, to avoid confusion with regard to the different parameters and their units used in our model, which enhances the reproducibility of our work, we transformed Eqs. S.1–S.3 so that they match the standard Volts and seconds scales. To do so, the coefficients and parameters in these equations had to be altered, which was done by assuming that $v$, $u'$ and $t'$ were not dimensionless at all but had units mV, mV/ms and ms instead. The dimensions of the terms in the right hand sides of the equations were matched to the left hand side:

$$\frac{dv}{dt} = 0.04 \frac{1}{\text{mV} \cdot \text{ms}} v^2 + 5 \frac{1}{\text{ms}} v + 140 \frac{\text{mV}}{\text{ms}} - u' + I' \tag{S.4}$$

$$\frac{du'}{dt} = a' \frac{1}{\text{ms}} (b' \frac{1}{\text{ms}} v - u') \tag{S.5}$$

$$\text{if } v \geq 30 \text{ mV, then } \begin{cases} v \leftarrow c' \text{ mV} \\ u' \leftarrow u' + d' \frac{\text{mV}}{\text{ms}} \end{cases} . \tag{S.6}$$

Transforming these equations to the standard Volts and seconds dimensions yielded

$$\frac{dV}{dt} = 40,000 \frac{1}{\text{V} \cdot \text{s}} V^2 + 5,000 \frac{1}{\text{s}} V + 140 \frac{\text{V}}{\text{s}} - u + I' \tag{S.7}$$

$$\frac{du}{dt} = a \frac{1}{\text{s}} (b \frac{1}{\text{s}} V - u) \tag{S.8}$$

$$\text{if } V \geq 0.030 \text{ V, then } \begin{cases} V \leftarrow c\text{V} \\ u \leftarrow u + d \frac{\text{V}}{\text{s}} \end{cases} . \tag{S.9}$$

The variables, parameters and coefficients in Eqs. S.7–S.9 have the appropriate standard scales that can be expressed in Volts and seconds. We may omit the unit notations and substitute $I' = \frac{I}{C_m}$ to obtain Eqs. 2–4 shown in the Method section of this article.

### Supplementary Figures

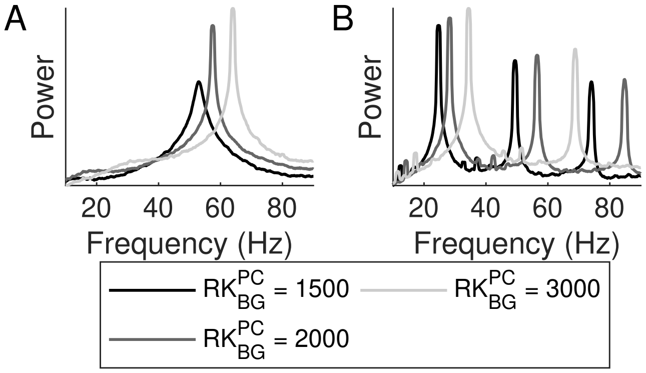


Figure S1: (A-B) Frequency spectra corresponding to peristimulus time histograms of the PCs when considering the pyramidal–PV cell circuit (A) and the pyramidal–SOM cell circuit (B) for various amounts of background input to the pyramidal cells ($RK_{bg}^{PC}$).

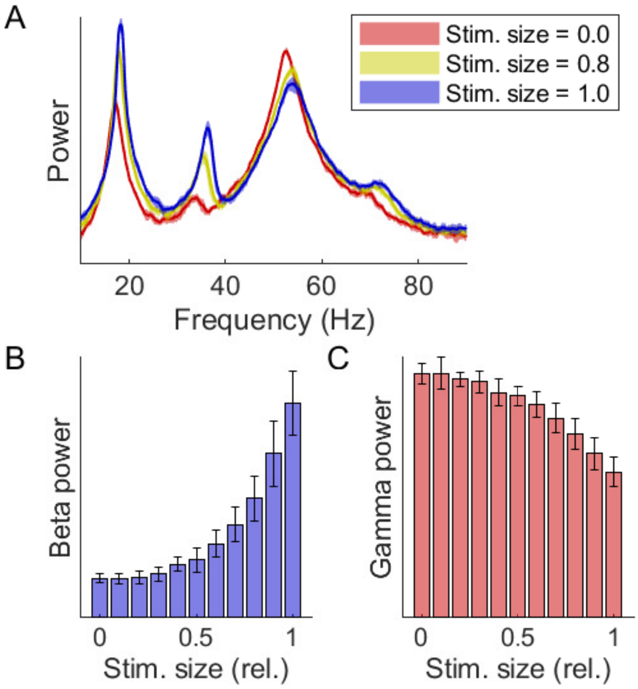


Figure S2: (A) Frequency spectra for three different relative stimulus sizes. Solid lines and shaded areas depict the *mean* ± *standard deviation* across repeats of the simulation with different random seeds. Spectra were obtained by considering the peristimulus time histograms of the PCs.
(B–C) Mean power across the beta (15–25 Hz, B) and the gamma (40–60 Hz, C) frequency bands as functions of the relative stimulus size. Bars and error bars depict the *mean* ± *standard deviation* across repeats of the simulation with different random seeds. Powers were derived from the spectra corresponding to the peristimulus time histograms of the PCs.
Abbreviations: rel. = relative, stim. = stimulus.

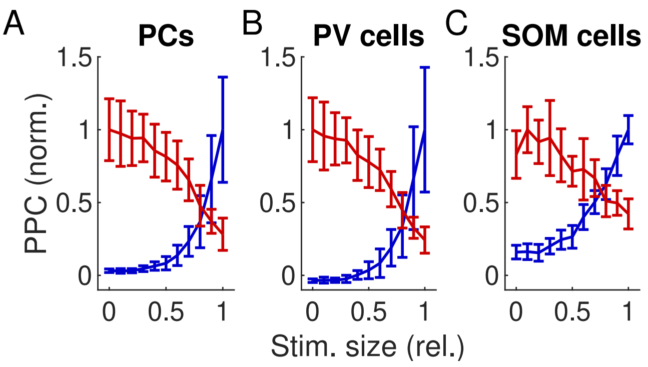


Figure S3: (A–C) Normalized spike-LFP pair-wise phase consistencies (PPCs) of the pyramidal (A), PV (B) and SOM (C) cells with respect to the beta ($15-25$ Hz) and gamma ($40-60$ Hz) frequency ranges as a function of the stimulus size. Red and blue correspond to the beta and gamma frequency band, respectively. Solid lines and error bars depict the *mean* $\pm$ *standard deviation* across repeats of the simulation with different random seeds. PPCs were determined on basis of the peristimulus time histograms of the PCS.
Abbreviations: coef. = coefficient, corr. = correlation, pyr. = pyramidal, rel. = relative, stim. = stimulus.

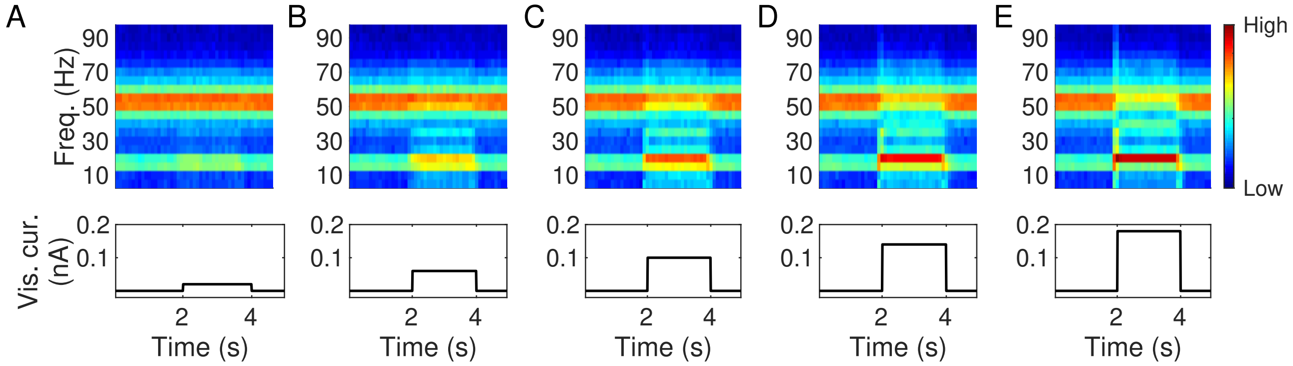


Figure S4: (A–E) Spectrograms of the PSTHs of the PCs (top row) and the concurrent change in the visual current ($I_{vis,0}$) throughout an epoch (bottom row). The latter plots show that this parameter was 0 pA in the first two and the final second of the epoch and could have a variety of values in the third and fourth second, namely 20 pA (A), 60 pA (B), 100 pA (C), 140 pA (D) or 180 pA (E). In the spectrograms, black and white colors correspond to a low and high power, respectively. The same color scale has been used for all spectrograms in this figure.

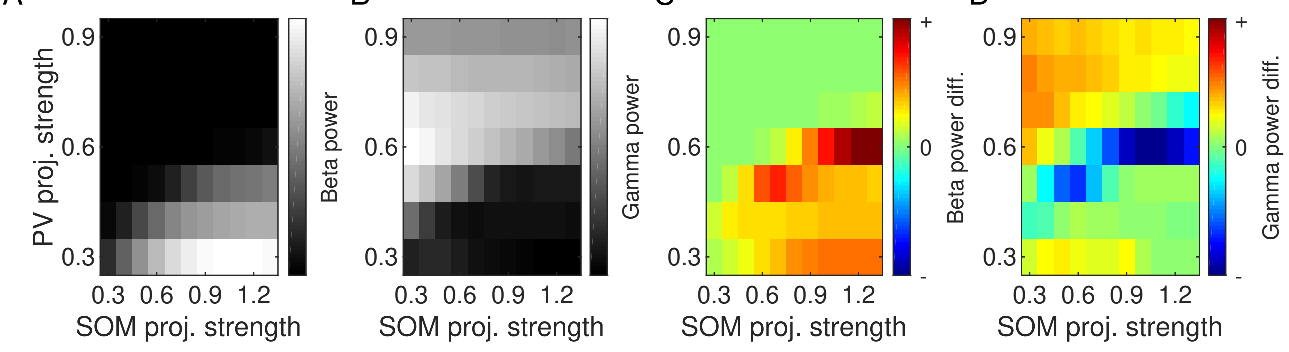


Figure S5: (A–B) Beta (A) and gamma (B) power as a function of the PV and SOM cell projection strengths. Also for these plots, none of the neurons received additional input through visually induced activity. Powers were derived from the frequency spectra obtained from the PSTHs of the PCs.
(C–D) Difference in beta (C) and gamma (D) power between the situation wherein the stimulated field covered the entire square patch and the one without any additional visual input. A positive difference means that the power is larger for visual stimulation. Powers were derived from the frequency spectra obtained from the PSTHs of the PCs.
Abbreviations: diff. = difference, proj. = projection.

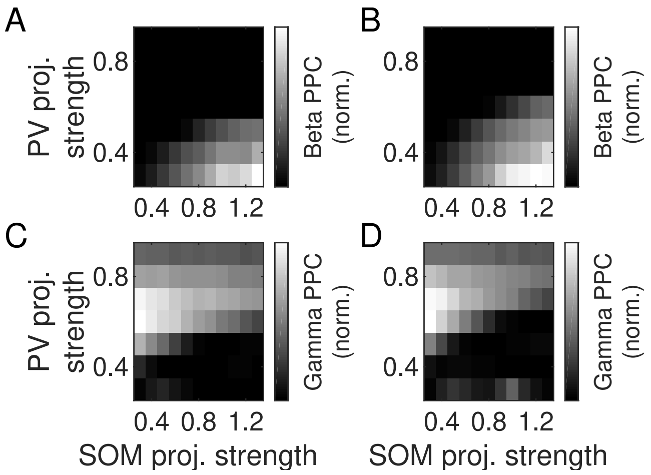


Figure S6: (A–D) Spike-LFP pair-wise phase consistencies (PPCs) for the rest state (left column) and the activated network (right column) with respect to the beta (A–B) and gamma (C–D) frequency bands. PPCs were obtained using the mean potentials of the PCs as LFP estimate.
Abbreviations: norm. = normalized, proj. = projection.

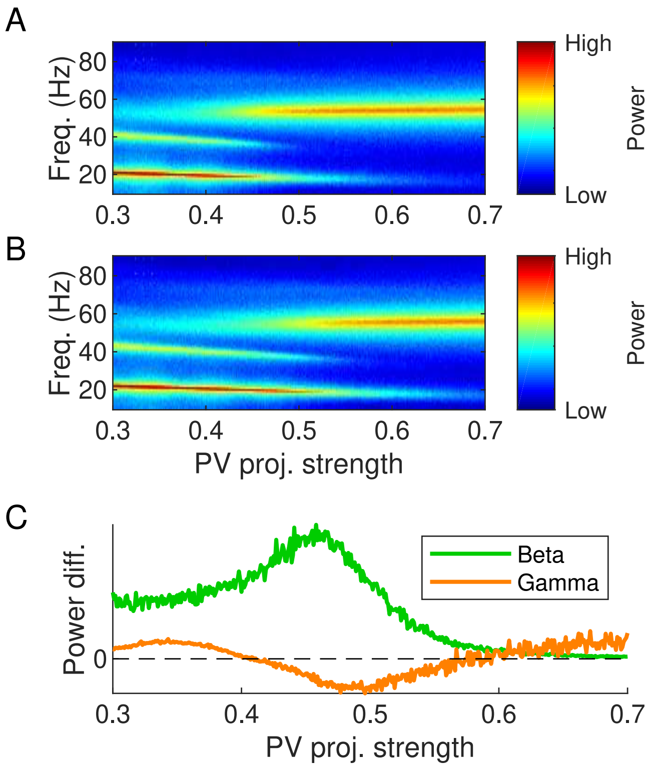


Figure S7: (A–B) Power density spectra of the resting-state ($D_{vis} = 0$, A) and the activated ($D_{vis} = 1$, B) network as a function of the PV cell projection strength for a fixed somatostatin expressing (SOM) cell projection strength ($S^{SOM} = 0.5$). The PSTHs of the PCs were used for the derivation of the spectra.
(C) Power differences between the activated and resting state network activity with respect to the beta (15–25 Hz, green) and gamma (40–60 Hz, orange) frequency band as a function of the PV cell projection strength for the same fixed SOM cell projection strength. A positive and negative difference reflect a higher and lower power in the activated situation, respectively. Powers were calculated on the basis of the PSTHs of the PCs.
Abbreviations: diff. = difference, freq. = frequency, proj. = projection.
